# VirtuousPocketome: A Computational Tool for Screening Protein-ligand Complexes to Identify Similar Binding Sites

**DOI:** 10.1101/2023.12.12.571207

**Authors:** Lorenzo Pallante, Marco Cannariato, Lampros Androutsos, Eric A. Zizzi, Agorakis Bompotas, Xhesika Hada, Gianvito Grasso, Athanasios Kalogeras, Seferina Mavroudi, Giacomo di Benedetto, Konstantinos Theofilatos, Marco A. Deriu

## Abstract

Protein residues within binding pockets play a critical role in determining the range of ligands that can interact with a protein, influencing its structure and function. Identifying structural similarities in proteins offers valuable insights into their function and activation mechanisms, aiding in predicting protein-ligand interactions, anticipating off-target effects, and facilitating the development of therapeutic agents. Numerous computational methods assessing global or local similarity in protein cavities have emerged, but their utilization is impeded by complexity, impractical automation for amino acid pattern searches, and an inability to evaluate the dynamics of scrutinized protein-ligand systems. Here, we present a general, automatic and unbiased computational pipeline, named *VirtuousPocketome*, aimed at screening huge databases of proteins for similar binding pockets starting from an interested protein-ligand complex. We demonstrate the pipeline’s potential by exploring a recently-solved human bitter taste receptor, i.e. the TAS2R46, complexed with strychnine. We pinpointed 145 proteins sharing similar binding sites compared to the analysed bitter taste receptor and the enrichment analysis highlighted the related biological processes, molecular functions and cellular components. This work represents the foundation for future studies aimed at understanding the effective role of tastants outside the gustatory system: this could pave the way towards the rationalization of the diet as a supplement to standard pharmacological treatments and the design of novel tastants-inspired compounds to target other proteins involved in specific diseases or disorders. The proposed pipeline is publicly accessible, can be applied to any protein-ligand complex, and could be expanded to screen any database of protein structures.

## Introduction

In the field of structural biology, it is widely recognized that there is a strong relationship between the three-dimensional structure of a protein and its function ^1^. The recognition and analysis of structural similarities in proteins can represent a valuable strategy to gain insights into protein functions. In particular, the comparison of protein binding sites represents a challenging area of interest in the field of biology and biochemistry to improve the understanding of protein-ligand interactions, predict off-target effects, facilitate the development of more selective and effective therapeutic agents, investigate drug repurposing strategies, and explore polypharmacological treatments^2^. In particular, drug repurposing, also known as drug repositioning or drug reprofiling, refers to the process of identifying new therapeutic uses for existing drugs that were originally developed for a different indication^3^. This approach offers several advantages, including potentially shorter development timelines, reduced costs, and a higher likelihood of success compared to developing entirely new compounds^4^. On the other hand, polypharmacology refers to the ability of a drug or a pharmacological agent to interact with multiple biological targets within an organism^5^. In other words, a polypharmacological drug can affect multiple pathways, receptors, or proteins simultaneously. This is in contrast to traditional drug development, where the focus is often on designing drugs that specifically target a single molecular target associated with a particular disease. Since several diseases involve multiple molecular targets, polypharmacology represents an interesting modality in terms of efficacy and adaptability to complex biological environments^6^.

The present work is inserted in this context and aims to develop a novel computational pipeline to pinpoint protein binding sites sharing similarities with an investigated target of interest. The identification of binding pockets with similar amino acid patterns can indicate potential off-target proteins for a given ligand, which can help in the design of drugs with minimal side effects. Additionally, the characterization of amino acid arrangements in binding pockets can contribute to the development of structure-based drug design methods to engineer drugs targeting specific protein families or selective ligands for individual proteins. This level of selectivity can be essential in the treatment of diseases, particularly when multiple proteins share similar functions but are involved in distinct physiological processes. As an example, we recently employed an embryonic version of the workflow presented here to understand the druggability of a query-binding site searching for similar motifs in proteins able to bind ligands of interest ^7^.

In the past years, several computational methods quantifying the global or local similarity of protein cavities have been developed^2,8^. In general, most of these methods share three main steps: (i) three-dimensional analysis of the structures of interest; (ii) structure comparison; and (iii) quantification of similarity through a metric (a scoring function). Many different representations of a given binding site are possible with varying degrees of retained information, e.g. the type of amino acid residues that interact with the ligand, or representing the binding site through a surface onto which the physical-chemical characteristics are projected, and even considering protein-ligand interactions. The first two methods can be regarded as structure-based, i.e. they stem from observing the structure of the protein. As far as the actual comparison strategies are concerned, these can be (a) graphical-theoretical approaches, where the maximum common subgraph is searched; (b) fingerprint approaches, where the shapes involved in the binding site are considered; (c) approaches based on labelled 3D points and geometric hashing, i.e. 3D transformations that align pairs of structures. Furthermore, comparison algorithms may or may not depend on the alignment of the structures of interest. Comparison methods that rely on residues can use graphs, fingerprints, or alternative approaches. In particular, the comparison reveals the similarity between the residues, the type of residues, and the atomic composition; also, such methods perform well where the sequence and atomic position of the structure of interest are well preserved. Those that rely on surfaces can instead use graphs or labelled 3D points for comparison. These methods are particularly used when dealing with binding sites in proteins that do not show significant conservation in residues, atomic composition, orientation, or folding, but show considerable selectivity towards common ligands. Indeed, in these cases, the distribution of the properties on the surface of the binding site and the shape of the binding site are determining factors for the selectivity of the ligands. And finally, methods that rely on interactions can use graphs or fingerprints for comparison ^9^. A summary of the main similarity search methods available in the literature is reported in Table 1 and a detailed comparison of the existing methodologies is available in recent litereature^9^.

**Table 1.**
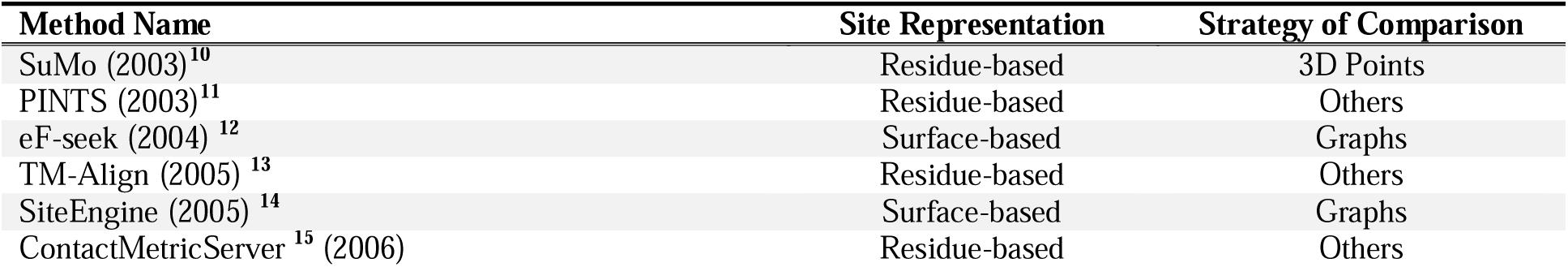
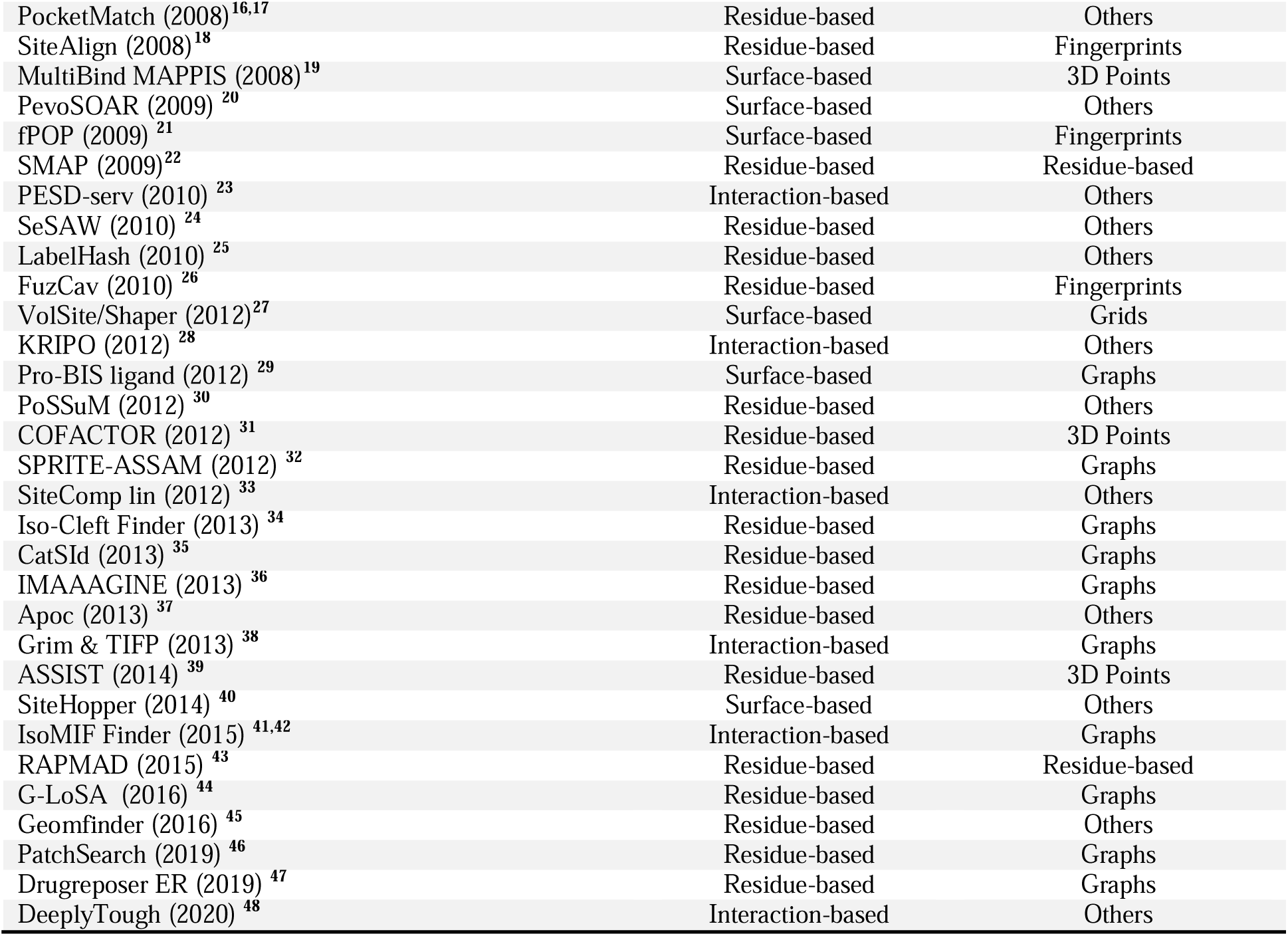
Summary of the main methods for the similarity search of a protein binding site in previous literature with their relative type of representation of the binding site and strategy of comparison.

The possibility of screening a high number of proteins for similar binding pockets can be particularly helpful and fruitful for those complex mechanisms and processes which may involve similar receptors and ligands for very different functions. In this context, we decided to turn our attention and use the proposed pipeline to explore some of the main actors involved in taste perception, due to the strong relationship between food intake and homeostasis regulation, disease onset, immune response and metabolism. The sense of taste is a sensory modality that plays a fundamental role in discriminating ingestible substances and nutrients from potentially harmful substances that must be avoided, especially in omnivorous species given the range of their feeding strategies ^49^. Humans, in particular, can perceive five primary taste qualities, i.e. sweet, umami, bitter, salty, and sour, through the interaction of molecules contained in food and specialized proteins, namely taste receptors, located on the papillae of the tongue. However, taste receptors are also expressed in other tissues besides the oral cavity, including the skin ^50,51^, brain ^52^, pancreas ^53,54^, heart ^55,56^, urethra ^57^, airway ^58^ and gastric ^59^ smooth muscle cells. Moreover, taste receptors are not only involved in the gustatory function, but they participate in other regulatory activities, such as regulation of metabolic activity ^60,61^, innate immune response and bronchodilatation ^60,62^, diabetes and obesity, glucose level maintenance, appetite regulation, as well as hormone release ^63^, and muscle contraction/relaxation ^64^.

In the present work, we decided to focus our attention on the bitter taste perception and relative actors, given the high scientific output produced in recent years concerning this specific taste sensation. Bitter taste receptors are the proteins responsible for the recognition of bitter foods, normally associated with potentially harmful substances. From a structural point of view, bitter taste receptors are GPCRs belonging to the taste 2 receptor family (*TAS2Rs*) ^65^ and are characterized by seven transmembrane helices (TMD) connected by three Intracellular Loops (ICLs) and three Extracellular Loops (ECLs)^66^. The structural core of the 7 TMD bundle is conserved across class-A GPCR and TAS2Rs. This core plays a fundamental role in the ligand binding in the extracellular (EC) region and information transduction in the intracellular (IC) region ^67^. Bitter taste receptors can be activated by a multitude of different agonists through various interaction types in their unique orthosteric binding pocket ^68,69^. Based on the chemical heterogeneity of their agonists, TAS2Rs have been distinguished into *promiscuous*, such as TAS2R10, TAS2R14, and TAS2R46, which are activated by a variety of chemically diverse compounds, and *selective*, activated instead by a limited number of similar compounds ^70^.

In the present work, we herein propose a novel, general and automatic algorithm, named VirtuousPocketome, to screen databases of protein structures to identify amino acid patterns that are similar to the ones forming the ligand binding site of a query receptor. Compared to previous literature, the novelty of this work resides in three main aspects: (i) the proposed pipeline accounts for the dynamics of the protein-ligand interaction by considering multiple binding site configurations obtained from a molecular dynamics (MD) trajectory; (ii) the identification of the crucial protein-ligand binding interactions is completely automatic; (iii) the results of the similarity search are filtered using an ad-hoc multi-step filtering process to exclude patterns unlikely to bind the ligand of interest. The algorithm builds structural motifs of the target binding site of the query receptor-ligand complex after clustering a molecular dynamics trajectory and then searches for similar patterns inside a specific protein database using the ASSAM code ^32^ followed by additional ad-hoc filtering steps. We applied our computational pipeline to screen the currently solved human proteome for proteins that exhibit a highly similar local amino acid pattern to the one lining the strychnine binding site in the human TAS2R46 bitter taste receptor. The rationale of the work is to explore the taste transduction pathway with a proteomic perspective to elucidate the possible role of tastants beyond the mere taste perception and to investigate whether other classes of proteins have a conserved ability to recognize such ligands, with possible implications in nutrition, homeostasis, and disease.

## Results

Details concerning convergence of the MD simulations and structural equilibrium of investigated molecular models are reported in the Supplementary Information (see also Figure S1 and Figure S2). The last 50 ns of each simulation replica were considered as structural equilibrium and were concatenated to obtain a final 150 ns-long trajectory representing the ensemble of protein conformations. These concatenated trajectories were used for the subsequent analysis and to search for similar binding pockets within the human proteome through the pipeline described herein.

### Motifs Creation

In the first step of the proposed pipeline, the motifs composed of the most important protein residues interacting with strychnine were identified. In particular, starting from the above-mentioned ensemble trajectory, residues within 10 Å from the position of the ligand have been extracted and their conformations clustered using the K-Means algorithm from MDAnalysis ^71^. Three clusters were identified as the optimum number through the silhouette method. After extracting the motif cluster centroids, we used PLIP to narrow down the most important non-covalent interactions which define the final three motifs. In all three of them, the ligand formed a salt-bridge interaction with residue GLU265^7.39^ and two hydrophobic interactions with residues TYR85^3.29^ and TRP88^3.32^, highlighting the stability of these contacts throughout the MD simulation. Furthermore, additional hydrophobic interactions were found with residues THR69^2.64^, PHE252^6.58^, and PHE261^7.35^ for motif 0 and with VAL61^2.56^, and VAL249^6.55^ for motif 2 (Figure 1). The three defined motifs were used for the subsequent similarity search step with ASSAM.

**Figure 1.**
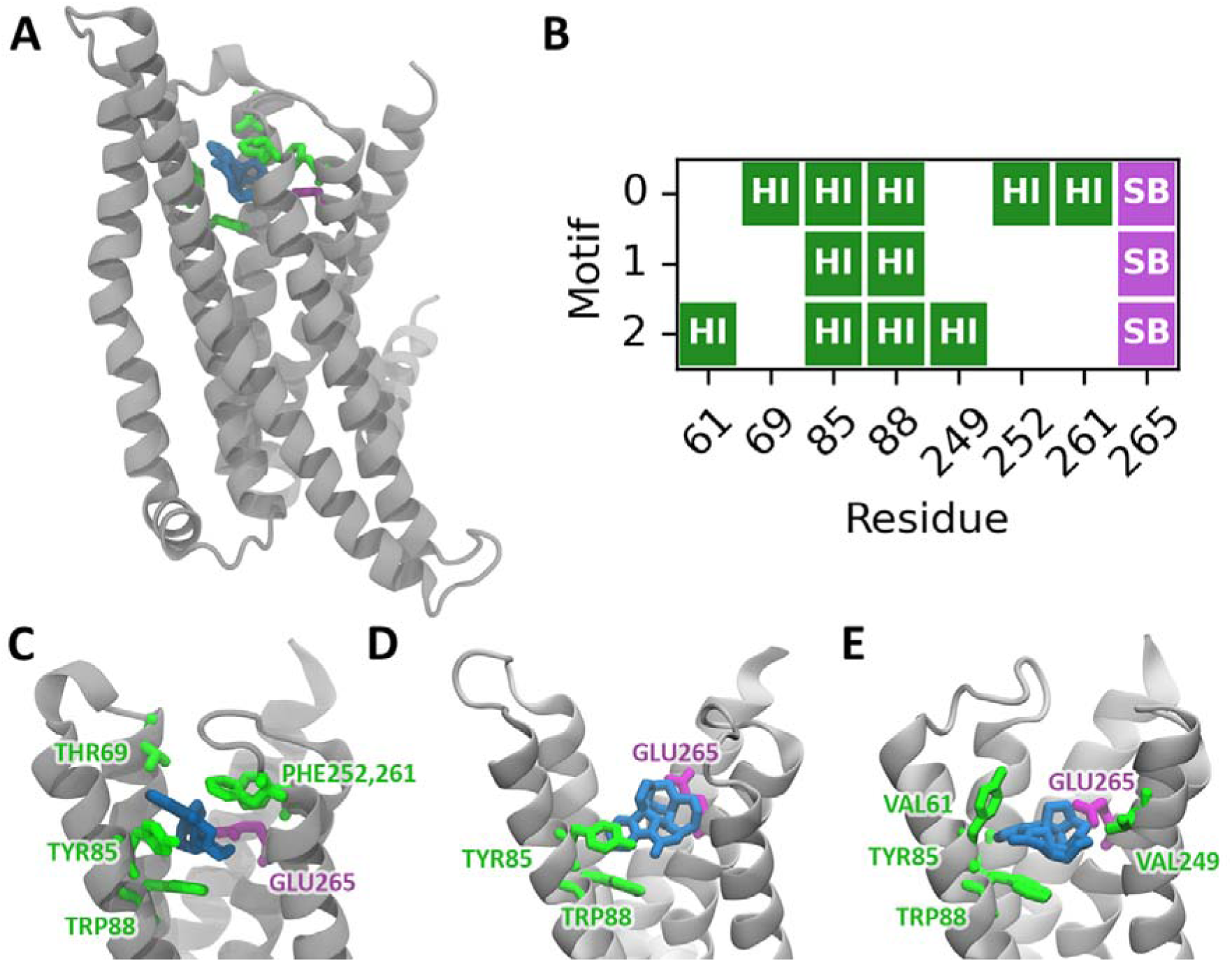
Main interactions defining the three motifs for the bitter taste receptor interacting with strychnine. (A) Representative snapshot of the bitter-ligand complex, (B) PLIP interaction analysis identifying Hydrophobic Interaction (HI) and Saltbridge Interaction (SB), (C, D, E) site views of the three motifs identified. The bitter taste receptor is represented in grey, the strychnine in blue and the interacting residues in green (hydrophobic interactions) and purple (salt bridges).

To check the importance of the identified residues, we also evaluated the *interaction probability* of protein residues with strychnine during the entire concatenated trajectory (500 ns x 3 replicas = 1500 ns). We calculated the interactions between the ligand and the receptor in each frame of the simulation (one frame every 200 ps) and we divided the total occurrences of a specific interaction by the total number of frames, obtaining an *interaction probability*. In line with the results on the cluster centroids, residues TYR85, TRP88, and GLU265 demonstrated the highest values with interaction probability higher than 80% (see also Table S1 and Figure S3). Interestingly, some weak interactions with the ligand in the initial experimental structure, such as those with residues ASN65 and THR180, are lost during the molecular dynamics simulation.

### Similarity Search and Multi-step Filtering

The similarity search against the entire human proteome consisting of 58972 structures (see the Database Curation section) resulted in a total of 6718 hits using the ASSAM code. The subsequent steps, i.e. the SASA and docking filtering steps, yielded a total of 1852 and 257 hits respectively (Figure 2A). In detail, we adopted a SASA threshold of 0.75, thus preserving all protein hits having a SASA in the binding pocket of at least 75% of the SASA of the original query binding site; on the other hand, we chose a docking threshold equal to 0.1, thus keeping the hits whose docking score would not differ by more than the 10% from the docking score of the query protein/ligand complex. The best hit in terms of docking score after the multi-step filtering process, namely PDB 3A4S, is represented in Figure 2B, highlighting the correspondence between the original motifs in the bitter taste receptor and the relative matching residues in the identified protein.

**Figure 2.**
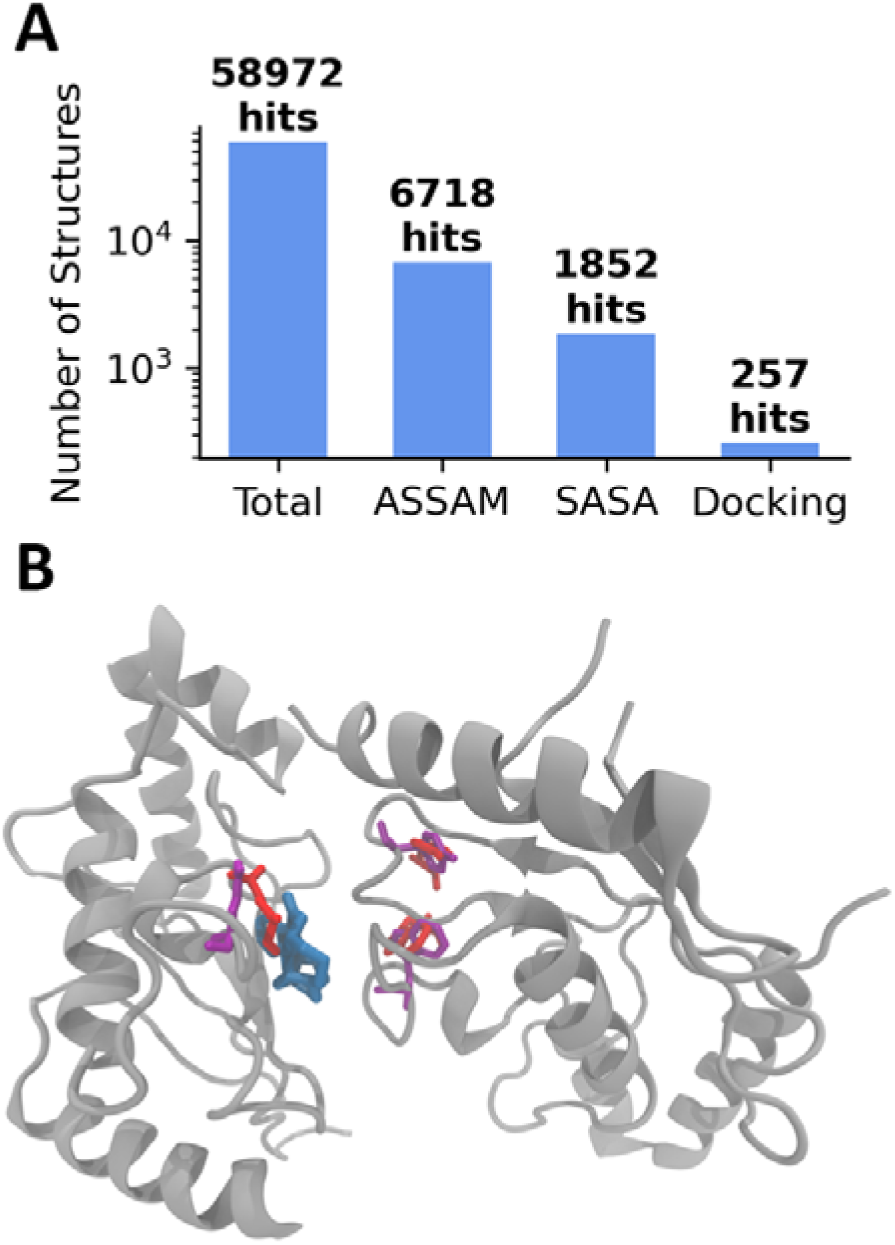
(A) Number of total structures in the original human proteome database and the number of selected hits from the similarity search and the subsequent multi-step filtering. (B) Binding site view of the strychnine bound to the best hit (PDB: 3A4S) according to the docking score at the end of the multi-step filtering process. Protein is rendered in grey, strychnine in blue, residues in the original motif of the bitter taste receptor in red and matching residues in the 3A4S structure in violet.

The pipeline automatically generated a report file (see ‘Results.txt’ file in the Supplementary Material) which includes the list of all the hits at the end of the screening process with additional information, such as PDB IDs with doi of the relative publication, protein classes, shared residues between hits and query, SASA and Docking Scores. The complete list of the 257 retrieved protein hits is reported in Table S2.

### Functional Enrichment and Signaling Pathway Analyses

VirtuousPocketome retrieved 145 unique Uniprot IDs relative to the previously identified hits, meaning that multiple PDBs at the end of the multistep filtering process corresponded to the same protein. The DAVID software was then employed to analyse the Gene Ontology terms and the signalling pathways data as described in the Methods section.

The functional enrichment analysis revealed that the input genes were significantly enriched for a total of 16 GO terms in the Cellular Components category, 10 terms in the Biological Processes category and 14 terms in the Molecular Functions category, based on the corrected p-value. In addition, 0 KEGG and 0 Reactome pathways were found to be significantly enriched at the same p-value threshold. The best 5 GO terms for each of the above-mentioned categories are represented in Figure 3, whereas all the significantly retrieved GO terms are represented in the Supplementary Information (Figure S4 and Figure S5). Regarding the Biological Processes (BP), most of the retrieved genes are related to metabolic processes (40.9%), including organic substance, cellular and nitrogen compound metabolic processes. Besides, the most represented Molecular Functions (MF) are related to the binding of different species, such as proteins, small compounds or ions, and to enzyme activity, such as transferase, hydrolase, and oxidoreductase. Finally, regarding the last analysed GO term, the most represented Cellular Components (CC) are cytoplasm and membrane (39.0%).

**Figure 3.**
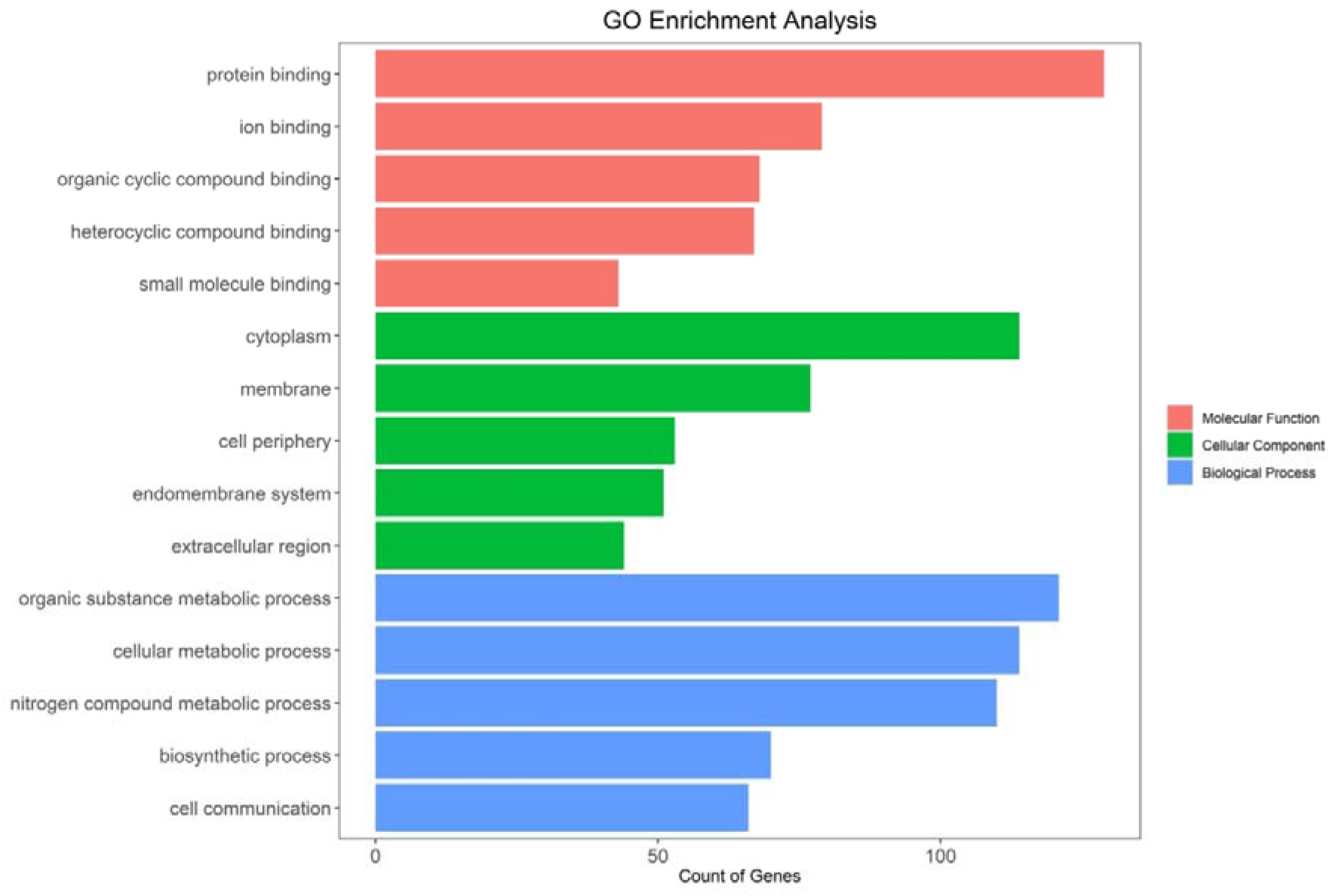
Bar plots representing the best 5 retrieved GO terms for each category in the third level of the GO hierarchy relative to Biological Processes (BP), Molecular Functions (MF) and Cellular Components (CC).

## Discussion

Herein, we developed a novel computational pipeline, named VirtuousPocketome, to screen a desired database for proteins sharing similar binding sites with a specific protein of interest. VirtuousPocketome is based on four major steps: (i) the motifs creation step, which identifies the most important residues involved in the interaction of the protein-ligand complex under investigation; (ii) the similarity search step, in which the human solved proteome is screened for similar motifs compared to the ones retrieved in the previous step; (iii) the multi-step filtering step, which preserves only the protein hits with binding sites effectively accessible and with a certain docking affinity for the query ligand; (iv) functional enrichment and signalling pathway analyses, which identifies the most relevant cellular components, molecular functions, biological processes and signalling pathways related to the protein hits identified in the previous steps. VirtuousPocketome takes as input the molecular structure (and eventually also the molecular dynamics trajectory) of a protein-ligand complex and gives as output a list of PDB structures sharing similar binding sites. The developed protocol is automatic and unbiased and can be applied to any protein-ligand complex.

To evaluate the developed code and prove its potential, we here selected the human TAS2R46 bitter taste receptor bound to a bitter ligand, i.e. strychnine and we screened the entire human proteome to search for similar structural motifs. The choice for this molecular system was driven by the recent experimental determination of the TAS2R46-strychnine complex structure ^72^, as well as the fact that strychnine is experimentally known to target not only TAS2R46 ^68^ but also other bitter taste receptors, such as TAS2R10 ^73^, and even other proteins ^74^. Moreover, bitter taste receptors have been widely investigated through computational molecular modelling in past years, ensuring enough data for comparison and evaluation of some of the results of the platform ^69,73,75–78^. Furthermore, the strong relationship between food intake and health status, including the regulation of homeostasis and metabolism, makes the chosen molecular machinery a particularly intriguing and relevant testbed for our computational screening pipeline to pinpoint possible secondary targets outside the gustatory system for food-related tastants.

The first step of the proposed pipeline, i.e. the motifs creation step, applied to the TAS2R46 bitter taste receptor bound to strychnine pinpointed three major motifs of residues mostly involved in the ligand binding. In particular, all three motifs shared a salt-bridge interaction with residue GLU265^7.39^ and two hydrophobic interactions with residues TYR85^3.29^ and TRP88^3.32^, whereas the first motif comprised also hydrophobic interactions with residues THR69^2.64^, PHE252^6.58^, and PHE261^7.35^ and the third motif with VAL61^2.56^, and VAL249^6.55^ (Figure 1). Interestingly, some of these residues have already been suggested by previous literature to be important interactions for ligand binding of bitter taste receptors. In detail, GLU265^7.39^ and TRP88^3.32^ were demonstrated to be pivotal in TAS2R46 activation by strychnine ^72^. Moreover, mutagenesis studies have reported the importance of residue GLU265^7.39^ for agonist responsiveness of TAS2R46 ^68^ and the same position has been also linked to ligand binding for similar receptors, including hydroxytryptamine (5-HT) receptors ^79,80^, adrenergic receptors ^81,82^, purinergic receptors ^83,84^, and cholecystokinin-B (CCK-B)/gastrin receptor ^85^. Moreover, residue TRP88^3.32^ is widely conserved among TAS2Rs and was found to be crucial in the activation of TAS2R43, TAS2R30 and TAS2R46 ^68,86^. The importance of residue in position 3.29 (TYR85 for TAS2R46) was confirmed also for TAS2R10, which shows a similar binding site to TAS2R46 and is also activated by strychnine ^73^. These pieces of the literature confirmed the reliability of the motifs creation step of the proposed pipeline to pinpoint the most important and relevant residues involved in the ligand binding. It is worth mentioning that the possibility of analysing a molecular dynamics trajectory allows the identification of multiple motifs, three in the present case, one for each identified cluster centroid, ensuring a more exhaustive sampling of the protein-ligand interactions.

Starting from the identified motifs, similar amino acid patterns were searched in the entire currently solved human proteome, consisting of 58972 structures. The similarity search using the ASSAM code resulted in 6718 hits. Then, the multi-step filtering reduced the number of detected sites to a total of 257 PDB structures, which represent 0.44% of the original database and 3.83% of the structures in ASSAM code output (Figure 2A). It is worth mentioning that in the final set of 257 hits notable entries include the potassium voltage-gated channel and acetylcholinesterase. The potassium voltage-gated channel is known to be modulated by strychnine (https://pubchem.ncbi.nlm.nih.gov/bioassay/588834). Although there is no experimental assay in the literature for the binding affinity between strychnine and acetylcholinesterase, strychnine is recognized to bind to the acetylcholine receptor, which binds acetylcholine similarly to the acetylcholinesterase^87,88^. The presented approach was revealed to be effective in discarding a high number of spurious proteins whose matching motifs were unlikely to bind strychnine, due to insufficient solvent exposure or low predicted affinity obtained from molecular docking. It is essential to highlight that while this pipeline can identify structures with amino acid patterns similar to a specified binding pocket, the actual ability of the ligand to interact with the identified proteins needs validation through experimental methods. This study is presented as a screening platform designed to pinpoint proteins that are more likely to be promising targets for the given ligand, with the understanding that experimental testing is crucial for confirmation.

The functional enrichment analysis allowed us to pinpoint the main biological processes, molecular functions, and cellular components related to the gene expressing the retrieved hit proteins at the end of the VirtuousPocketome pipeline (Figure 3). Most of the biological processes highlighted are connected to metabolic processes, which seems an intriguing result considering the strong relationship between taste perception, food intake and metabolism. Indeed, these results might indicate that strychnine has not only the ability to activate the TAS2R46 bitter taste receptor and elicit the bitter taste sensation, but it should be also the potential trigger or modulator of proteins directly involved in the metabolic processes. Moreover, the protein hits are mainly involved in molecular functions related to the binding of proteins or other small compounds and are localized mainly in the cytoplasm and membrane. These results seem pretty reasonable considering that TAS2R46 is a transmembrane protein and a promiscuous bitter taste receptor, able to bind a wide spectrum of chemical compounds. In light of these results, it is worth mentioning that strychnine is known to bind glycine and acetylcholine receptors ^74^, which are membrane proteins involved in transport and signalling functions. Therefore, the presented pipeline was able to detect similar binding sites in receptors with the same localization and biological function of known strychnine targets. In addition, strychnine is a rather promiscuous ligand with anti-plasmodial and anti-cancer activity and other yet-unresolved molecular targets ^89^.

In conclusion, in the present work, we developed a novel, general and automatic pipeline for identifying proteins sharing similar binding pockets with a query receptor-ligand complex by screening the entire solved human proteome. The developed pipeline is also freely accessible at https://virtuous.isi.gr/#/virtuous-pocketome and will be easily expanded in the future to other protein databases. We used the TAS2R46-strychnine complex as a testbed for the proposed method to investigate if other proteins except the taste receptors might share similar binding pockets for the recognition of tastants. The proposed methodology allows for a deep investigation of the TAS2R46-specific residues needed for the strychnine binding to pinpoint other human proteins sharing similar binding pockets and investigate the potential roles that this tastant could play in contexts beyond the gustatory system. The retrieved proteins could be further analyzed to predict whether the interaction with the compound of interest results in their activation or modulation. This will increase our understanding of possible secondary effects of tastants beyond the mere taste perception and their impact on the biological processes and molecular functions in which the retrieved hit proteins are involved. This approach can also assist the design of specific foods and ingredients to develop personalised treatments able to target desired proteins or receptors involved in specific processes or diseases with the ultimate goal of accessing the potential of the diet as a supplement to traditional pharmacological treatments.

## Methods

### VirtuousPocketome Workflow

#### Overall Workflow

The proposed computational pipeline requires as mandatory inputs the coordinates (PDB file) of a protein-ligand complex and the chain label(s) uniquely identifying the receptor(s) and the ligand in the provided structure. In order to allow the ligand to accommodate in the binding site and to assess the dynamic nature of the protein-ligand binding, the user can specify a molecular dynamics trajectory (GROMACS xtc/trr/pdb file), thus considering several configurations of the investigated complex. Moreover, additional custom parameters can also be defined by experienced users if needed (details in the following).

At present, the code is limited to the screening of the entire human proteome, but it can be easily expanded to any PDB-like database.

The overall workflow of the algorithm designed in the present work is divided into four main steps, each of which will be described in detail in the following paragraphs: ***Motifs Creation, Similarity Search, Multi-step Filtering, and Enrichment and Signaling Pathway Analyses*.**

The main output consists of a Txt file collecting the PDBs of the identified proteins sharing similar accessible binding site(s) compared to the query receptor-ligand complex. Additionally, the code provides as output the list of unique UniProt IDs related to the retrieved proteins, plots summarising the main results (see the Results section), and visualisation states of the molecular system under investigation.

The overall workflow is represented in Figure 4 in a flow chart representation.

**Figure 4.**
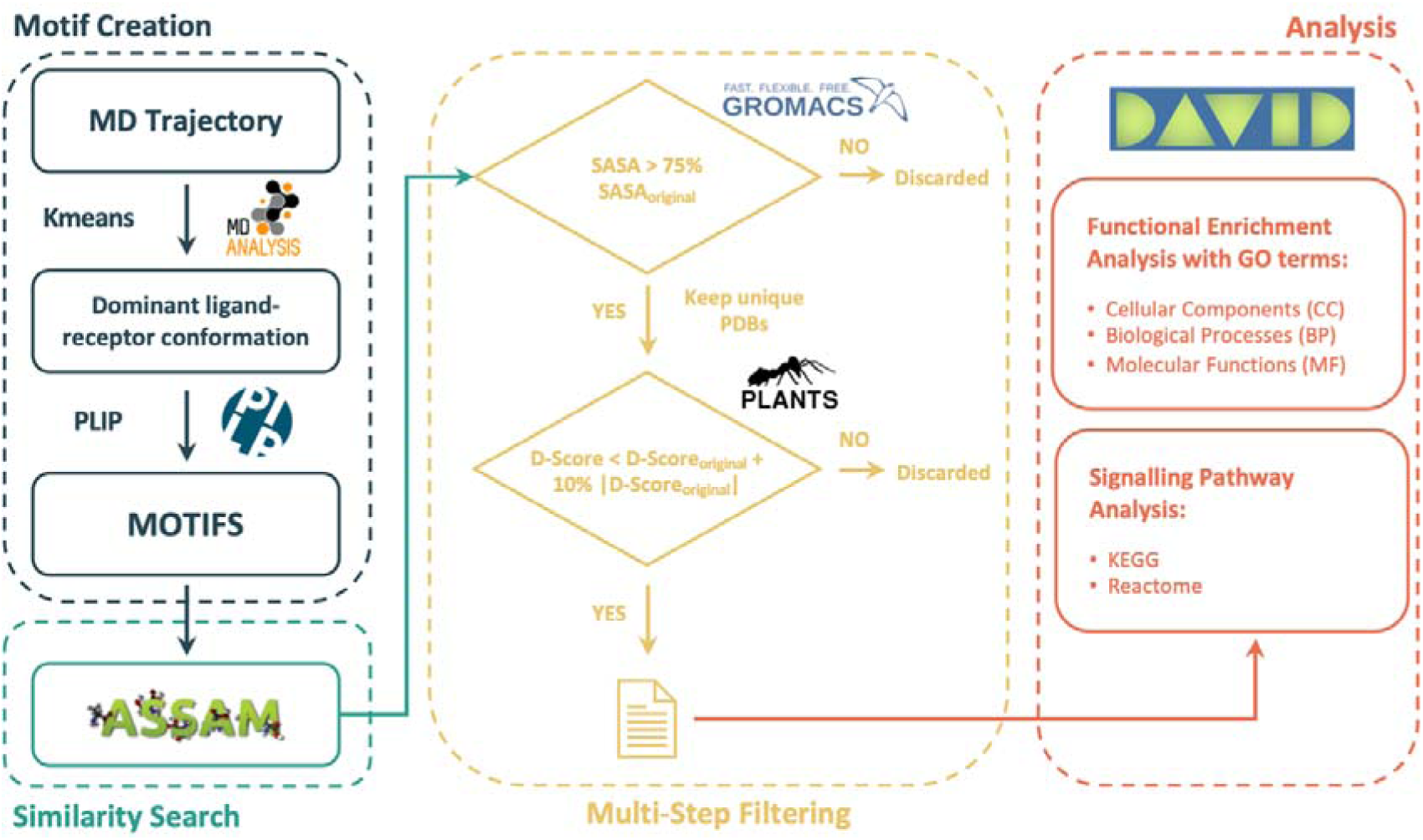
Flow Chart of the overall workflow of VirtuousPocketome.

For the implementation of this workflow, we mainly used GROMACS functionalities ^90^, MDAnalysis modules ^71^, and pdb-tools ^91^. The ASSAM source code was kindly provided by Prof. M. Firdaus Raih from the Molecular Function Regulation Lab (http://mfrlab.org).

#### Step 1 - Motifs Creation

Starting from the provided PDB input file and the MD simulation (if present) of the protein-ligand complex, a list of residues defining the binding site is retrieved based on the distance between the ligand and the receptor. The default distance threshold is set to 10 Å, but this can be overridden with a custom value provided by the user as an additional parameter. If the user only provides a single PDB file of the protein-ligand complex, a single binding site is defined according to the chosen distance threshold between the ligand and the protein. Conversely, if the user provides also the MD trajectory, the obtained subset of coordinates from each frame of the simulation identifying the ligand and the residues of the protein binding site are clusterized with the K-Means algorithms implemented in MDAnalysis, which is based on the distances between atoms positions ^71^. The user can set the desired number of clusters (k), otherwise, the optimal number of clusters is retrieved using the silhouette method ranging from a minimum of 2 clusters up to a maximum of 12 clusters. The silhouette method, as provided by the built-in scikit-learn function (sklearn.metrics.silhouette_score), is an example of an evaluation metric to indicate if clusters are well-defined. This method computes the mean distance between a sample and all other points within the same class (a) and the mean distance between a sample and all other points in the nearest neighbouring cluster (b). The Silhouette Coefficient, s, for a single sample, is then calculated as reported in Equation 1:

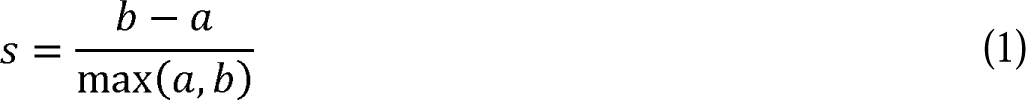

Then, the total Silhouette Coefficient is calculated as the mean of the Silhouette Coefficient for each sample. A higher Silhouette Coefficient score is indicative of denser and well-separated clusters, aligning with the conventional understanding of a cluster. It suggests that the samples within each cluster are tightly packed and distinct from samples in other clusters. This reflects a higher degree of cohesion within clusters and a greater separation between them. Therefore, the algorithm calculates the silhouette coefficient for different numbers of clusters (from 2 to 12) and then chooses the optimal number of clusters according to the best-achieved silhouette coefficient score. The subsequent centroids of the clusters are saved.

Starting from the previously defined binding sites, the subset of residues forming the binding site is further refined, highlighting those residues involved in non-covalent interactions with the ligand. These interacting residues are retrieved using the Protein-Ligand Interaction Profiler (PLIP) software ^92^, which detects hydrogen bonds, hydrophobic contacts, pi-stacking, pi-cation interactions, salt bridges, water bridges, metal complexes, and halogen bonds between ligands and targets. This two-step analysis allows the definition of the most relevant protein residues for the interaction with the ligand. This subset of residues will be defined as *motifs* in the following. If only the PDB file is passed by the user, a single motif will be created; if an MD trajectory is provided, a motif will be produced for each centroid of the binding site.

#### Step 2 - Similarity Search

In the second step, the similarity search for each extracted motif is carried out using the ASSAM software. The fundamental principles underlying the search methodologies of ASSAM are described in previous literature ^32,93^. In brief, the protein structure is represented as a graph, where nodes represent individual amino acid side chains and the geometric relationships between nodes form the edges of the graph. Each node is composed of two pseudo-atoms, which generate vectors that correspond to the nodes in the graph. The positions of these pseudo-atoms are strategically chosen to emphasize the functional portion of the corresponding side chain. The geometric relationships between pairs of residues are defined by distances calculated between their corresponding vectors, and these relationships are represented as the edges of the graph. If we let S, M and E denote the start, middle and end of a vector, the edges of the graph encompass five components: SS, SE, ES, EE, and MM distances, although only a subset of these distances is typically used to specify a query pattern. ASSAM employs a maximal common subgraph (MCS) approach using the Bron and Kerbosch MCS algorithm to enumerate all possible correspondences with similar protein patterns ^94^.

The choice to utilize ASSAM was made after an extensive examination of tools documented in the literature for assessing protein binding pockets (see also Table 1). Specifically, we scrutinized various methods, emphasizing those deemed most suitable for our objectives and compatible with integration into our pipeline. We specifically looked for methods that could compare a particular binding site with a broad set of proteins in a database (customizable if necessary), be executable locally rather than solely as a web server, provide output data on similar residues between the query protein and the target protein, perform searches quickly, and not require the prior definition of binding sites for proteins in the search database. This analysis pointed ASSAM as the most appropriate for the present work. In particular, ASSAM was chosen for (i) the simplicity in defining the binding site to be searched, i.e. a coordinate file containing the residues of interest in PDB format, (ii) the possibility to screen against any desired database in PDB format, (iii) the comparably fast time to solution, and (iv) the possibility of considering and preserving both right-handed and left-handed orientations of the alpha-helices to retrieve the superpositions. A left-handed α-helical bundle superimposed onto a right-handed one is not equivalent, particularly concerning the handedness of the two groupings of amino acids. However, when it comes to chemical activity, two groupings of amino acids may exhibit the same behaviour despite having different handedness. The crucial factor, in this case, is the distance between the individual residues ^32^. Thanks to the source code kindly provided by the ASSAM developers, the proposed pipeline does not rely on the ASSAM web-based analysis tools and runs entirely offline on-premises.

The output of the ASSAM search is formatted as a list, in which each row corresponds to a *hit* protein found in the screened protein database, along with its PDB accession ID, which presents a match with the residues of the input motif, also referred to as *query*. The matching residues between the query and the hit are also indicated, as well as the root-mean-square deviation (RMSD) value between query residues and hit residues after alignment. Finally, additional pieces of information are retrieved, namely the number of the initial conformation from which the query motif was created, and further information on the hit protein obtained from the RCSB Protein Data Bank site, such as the DOI of the corresponding publication and the EC classification.

#### Step 3 - Multi-step Filtering

To further refine the output from the previous steps and select only the protein hits with accessible and high-affinity binding sites, two additional filtering steps, i.e. (i) the SASA and (ii) the docking filters, have been implemented.

The first step involves the calculation of the Solvent-Accessible Surface Area (SASA) using GROMACS ^90^. In detail, for each hit protein, the SASA, evaluated in nm^2^, is calculated by measuring the value for all the residues matching the residues of the query protein’s motifs. Values equal to zero indicate that the cleft identified by the residues is not solvent-exposed, but is rather buried in the structure of the protein and therefore likely inaccessible for the solvent or the ligand. Values that are greater than zero, on the other hand, indicate that the cleft formed by the hit residues is on the surface of the corresponding protein and therefore might allow for ligand binding. Only hits with the binding site having SASA greater than a predefined threshold of the SASA value of the corresponding query motif are retained. The SASA criterion is summarised by equation (1):

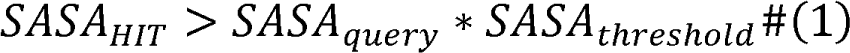

Where SASA_HIT_ corresponds to the calculated SASA value of the given hit motif, SASA_query_ is the SASA value of the corresponding motif of the query protein-ligand complex, and SASA_threshold_ is the threshold to select only hits with the desired SASA. SASA_threshold_ is set to 75% by default, but the user can specify a custom value in the additional input parameters. The selection of this threshold was imposed as a compromise to obtain a reasonable number of protein hits that could be effectively and rationally considered while retaining targets with motifs that are solvent-exposed and prone to ligand docking. The decision to set as default a high SASA threshold is based on the importance of solvent-accessible surface area (SASA) in assessing the capability of a protein binding site to accommodate ligands. Considering only SASA values considerably lower than the original protein binding site could compromise the algorithm’s ability to retain only those binding sites most similar to the reference complex and thus more prone to ligand binding. Setting the SASA threshold to higher values, on the other hand, may lead to the exclusion of potentially promising binding sites and protein hits. As a result, users have the flexibility to adjust this threshold based on their screening objectives.

The second filtering step relies on the molecular docking of the original ligand in the query complex onto the retrieved hits from the previous steps. The docking procedure was implemented using SPORES and PLANTS ^95,96^. PLANTS software was chosen since it exhibited excellent performance in terms of pose prediction and time to solution compared to other molecular docking software ^97,98^ and since it has been already successfully used for molecular docking and virtual screening campaigns on GPCRs ^99–103^. These qualities make PLANTS particularly well-suited for the present work. Only hits with binding sites exhibiting a docking score (DSCORE) similar to or below the docking score of the original protein-ligand complex are retained. In particular, the docking score of the selected hit (DSCORE_HIT_) is compared with the docking score on the original query complex 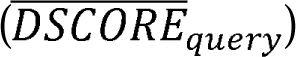 adjusted by a defined percentage using the specified docking threshold (DSCORE_threshold_). The docking threshold is set by default as 10% of the original protein-ligand complex docking score, if only a PDB is provided as input, or the 10% of the average PLANTS docking scores between the centroids of the original protein-ligand complex, if the MD trajectory was specified. The docking filtering criterion is therefore defined by equation (2):

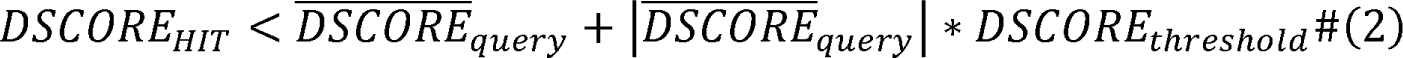

#### Step 4 - Enrichment and Signalling Pathway Analyses

In this step, functional enrichment and signalling pathway analyses were performed to collect information regarding roles, functions, distributions, expressions and pathways in which the identified targets from the previous steps are involved. The pipeline automatically retrieves the unique UniProt IDs of the protein hits (since several PDB codes can correspond to the same protein) and searches for related genes using the DAVID functional annotation program ^104,105^. We decided to focus our attention on the Gene Ontology (GO) terms ^106^ related to the Cellular Components (CC), Biological Processes (BP) and Molecular Functions (MF). The resulting GO terms were used to identify significantly enriched functional categories, using a statistical approach that takes into account the size of the protein list, the number of genes associated with each GO term, and the background gene set. We also performed pathway analysis using the Kyoto Encyclopedia of Genes and Genomes (KEGG) and the Reactome databases to identify the most represented pathways ^107–109^. GO terms and KEGG pathways with corrected for multiple testing q-value < 0.1 were considered to be significantly enriched. The correction of the p-values and the calculation of the q-values were performed using the Benjamini-Hochberg FDR adjustment method ^110^. The pipeline automatically generates separate plots with the above-mentioned analysis.

#### Database Curation

The developed tool can screen a database of target proteins in the PDB format for binding pocket similarities against a query protein-ligand complex. For the present work, we have collected and refined all the experimentally solved PDB structures belonging to the human proteome. We first retrieved all the solved PDB codes belonging to *homo sapiens* from the NCBI database ^111^ using a dedicated Python API (Bio.Entrez package), and obtained a total of 60159 entries (1^st^ February 2023). We downloaded all the found queries from the RCSB database (http://www.rcsb.org/) ^112^, reaching a total of 59267 structures (892 PDBs were not available). Then, the database was cleaned by removing (i) the chains in the PDBs not belonging to the human organism (such as in the case of protein chimaeras), (ii) multiple models from each PDB, (iii) ANISOU and HETATM lines, (iv) alternative locations of the atoms in the PDB by preserving the ones with the highest occupancy. At the end of this cleaning protocol, we ended up with 58972 PDB files.

### Molecular Modelling and Dynamics

#### System Setup

The previously described workflow was applied to search for proteins sharing similar binding pockets with a human bitter taste receptor. We employed the recently-solved TAS2R46 human bitter taste receptor bound with an agonist bitter compound, named strychnine (PDB ID: 7XP6) ^72^. We first removed undesired molecules and structures from the PDB, preserving only the bitter taste receptor and strychnine. Since the receptor structure presents some missing residues (157-172), we downloaded the relative model (sequence P59540 of Homo Sapiens hTAS2R46) from the AlphaFold Protein Structure Database ^113^. The AlphaFold model was then aligned to the receptor in the 7XP6 PDB. Missing residues in the original experimental structure were then built by homology modelling with MOE (Molecular Operating Environment) software ^114^ using the relative portion of the AlphaFold model as a template, thus filling the gap in the original experimental structure. Then, the structure of strychnine was refined using MOE, assigning the correct protonation at neutral pH and salt concentration of 0.15 M (see also Figure S6). As reported in previous literature^72^, the tertiary amine of the molecule is protonated and its total charge is +1, as also reported in the DrugBank database (https://go.drugbank.com/drugs/DB15954).

The obtained complex was embedded into a POPC membrane using the CHARMM-GUI web server ^115^. The receptor was aligned with the membrane bilayer using the OPM web server included in the CHARMM-GUI preparation protocol. The final number of POPC lipids (165) was set by a trial-and-error approach to creating a box large enough to meet the minimum image convention for the protein during the MD simulation: the final box size was set to 8.36 x 8.36 x 11.47 nm, with the z direction perpendicular to the cell membrane plane. The system box was filled with water and neutralized using NA^+^ and CL^-^ ions at 0.15 M physiological concentration.

We used the AMBER forcefield to describe the molecular system: in detail, we used the Lipid-21 forcefield ^116^ for lipids, the AMBER19SB ^117^ for the protein, ions and water and the General Amber Force Field (GAFF2) forcefield ^118^ to obtain the topology for strychnine, as implemented directly in the CHARMM-GUI suite.

#### Simulation Protocol

The simulation workflow suggested by CHARMM-GUI was followed during minimisation, equilibration with position restraints and final simulation production. First, the system was energy-minimized using the steepest descent algorithm for 5000 steps. Then, six equilibration steps were performed gradually reducing the position restraints on the lipids and protein-heavy atoms (from 1000 to 0 kJmol^-1^nm^-1^ for lipids, from 4000 to 50 kJmol^-1^nm^-1^ for the protein backbone and from 2000 to 0 kJmol^-1^nm^-1^ for protein side-chain heavy atoms). The system was equilibrated in the NVT ensemble for 250 ps with a conservative timestep of 1 fs, using the Berendsen thermostat ^119^ with a coupling time constant of 1 ps and a reference temperature of 303.15 K, which is above the phase-transition temperature for POPC, and subsequently in the NPT ensemble for 125 ps with the same 1 fs timestep, followed by a further simulation of 375 ps with a 2 fs timestep, using the Berendsen thermostat with the same parameters as before and the Berendsen barostat ^119^ with semi-isotropic pressure coupling at 1 atm with a coupling time constant of 5 ps. Overall, the systems underwent 750 ps of equilibration.

The TAS2R46 bitter taste receptor system was then simulated for 400 ns without position restraints with a 2 fs time step in the NPT ensemble. Temperature coupling was done with the Nose-Hoover algorithm ^120^, whereas, pressure coupling with the Parrinello-Rahman algorithm ^121^. The PME algorithm was used for electrostatic interactions with a cut-off of 0.9 nm. A reciprocal grid of 72 x 72 x 96 cells was used with 4th-order B-spline interpolation. A single cut-off of 0.9 nm was used for Van der Waals interactions. LINCS (LINear Constraint Solver) algorithm for h-bonds ^122^ was applied in each simulation step.

Three simulation replicas were performed to ensure the reproducibility of the simulations and to enlarge the simulation statics. All Molecular Dynamics (MD) simulations were performed using GROMACS ^90^ and the Visual Molecular Dynamics (VMD) package was employed for the visual inspection of the simulated systems ^123^.

## Acknowledgements

The present work has been developed as part of the VIRTUOUS project, funded by the European Union’s Horizon 2020 research and innovation program under the Marie Sklodowska-Curie-RISE Grant Agreement No. 872181 (https://www.virtuoush2020.com/).

The authors would also like to acknowledge Prof. M. Firdaus Raih from the Molecular Function Regulation Lab (http://mfrlab.org) for kindly providing us with the source code of ASSAM software.

## Author Contributions

**Lorenzo Pallante**: Conceptualization; Data curation; Formal analysis; Methodology; Software; Validation; Visualization; Writing - original draft; Writing - review & editing.

**Marco Cannariato**: Conceptualization; Data curation; Formal analysis; Methodology; Software; Validation; Visualization; Writing - original draft; Writing - review & editing.

**Lampros Androutsos**: Conceptualization; Data curation; Formal analysis; Investigation; Validation; Visualization; Writing - review & editing.

**Eric A. Zizzi**: Conceptualization; Supervision; Methodology; Writing - review & editing;

**Agorakis Bompotas**: Supervision; Software; Methodology; Visualization; Writing - review & editing;

**Xhesika Hada:** Conceptualization; Methodology; Writing - review & editing;

**Gianvito Grasso:** Supervision; Writing - review & editing;

**Athanasios Kalogeras:** Supervision; Writing - review & editing;

**Seferina Mavroudi:** Supervision; Writing - review & editing;

**Giacomo Di Benedetto:** Supervision; Writing - review & editing;

**Konstantinos Theofilatos:** Conceptualization; Supervision; Formal analysis; Validation; Writing - review & editing;

**Marco Agostino Deriu**: Conceptualization; Formal analysis; Funding acquisition; Project administration; Resources; Supervision; Validation; Visualization; Writing - review & editing.

## Data Availability

All data to replicate the simulations are accessible at https://github.com/M3B-Lab/VirtuousPocketome.

## References

1. Artymiuk, P. J., Poirrette, A. R., Rice, D. W. & Willett, P. A polymerase I palm in adenylyl cyclase? Nature 388, 33–34 (1997).

2. Ehrt, C., Brinkjost, T. & Koch, O. A benchmark driven guide to binding site comparison: An exhaustive evaluation using tailor-made data sets (ProSPECCTs). PLoS Comput Biol 14, e1006483 (2018).

3. Hodos, R. A., Kidd, B. A., Shameer, K., Readhead, B. P. & Dudley, J. T. *In silico* methods for drug repurposing and pharmacology. WIREs Mechanisms of Disease 8, 186–210 (2016).

4. Independent Interdisciplinary Consultant, Oxford, United Kingdom & Flower, D. Drug Discovery: Today and Tomorrow. Bioinformation 16, 1–3 (2020).

5. Kabir, A. & Muth, A. Polypharmacology: The science of multi-targeting molecules. Pharmacological Research 176, 106055 (2022).

6. Chaudhari, R., Fong, L. W., Tan, Z., Huang, B. & Zhang, S. An up-to-date overview of computational polypharmacology in modern drug discovery. Expert Opinion on Drug Discovery 15, 1025–1044 (2020).

7. Cannariato, M., Miceli, M. & Deriu, M. A. In silico investigation of Alsin RLD conformational dynamics and phosphoinositides binding mechanism. PLOS ONE 17, e0270955 (2022).

8. Eguida, M. & Rognan, D. Estimating the Similarity between Protein Pockets. IJMS 23, 12462 (2022).

9. Ehrt, C., Brinkjost, T. & Koch, O. A benchmark driven guide to binding site comparison: An exhaustive evaluation using tailor-made data sets (ProSPECCTs). PLoS Comput Biol 14, e1006483 (2018).

10. Jambon, M., Imberty, A., Deléage, G. & Geourjon, C. A new bioinformatic approach to detect common 3D sites in protein structures: Detection of 3D Sites in Protein Structures. Proteins 52, 137–145 (2003).

11. Stark, A., Sunyaev, S. & Russell, R. B. A Model for Statistical Significance of Local Similarities in Structure. Journal of Molecular Biology 326, 1307–1316 (2003).

12. Kinoshita, K. Identification of Protein Biochemical Function by Searching the Similar Shape and Electrostatic Potential on the Molecular Surface of Proteins. Seibutsu Butsuri 44, 150–154 (2004).

13. Zhang, Y. TM-align: a protein structure alignment algorithm based on the TM-score. Nucleic Acids Research 33, 2302–2309 (2005).

14. Shulman-Peleg, A., Nussinov, R. & Wolfson, H. J. SiteEngines: recognition and comparison of binding sites and protein-protein interfaces. Nucleic Acids Research 33, W337–W341 (2005).

15. Lisewski, A. M. & Lichtarge, O. Rapid detection of similarity in protein structure and function through contact metric distances. Nucleic Acids Research 34, e152–e152 (2006).

16. Nagarajan, D. & Chandra, N. PocketMatch (version 2.0): A parallel algorithm for the detection of structural similarities between protein ligand binding-sites. in 2013 National Conference on Parallel Computing Technologies (PARCOMPTECH) 1–6 (IEEE, Bangalore, India, 2013). doi:10.1109/ParCompTech.2013.6621397.

17. Yeturu, K. & Chandra, N. PocketMatch: A new algorithm to compare binding sites in protein structures. BMC Bioinformatics 9, 543 (2008).

18. Schalon, C., Surgand, J., Kellenberger, E. & Rognan, D. A simple and fuzzy method to align and compare druggable ligand[binding sites. Proteins 71, 1755–1778 (2008).

19. Shulman-Peleg, A., Shatsky, M., Nussinov, R. & Wolfson, H. J. MultiBind and MAPPIS: webservers for multiple alignment of protein 3D-binding sites and their interactions. Nucleic Acids Research 36, W260–W264 (2008).

20. Tseng, Y. Y., Dundas, J. & Liang, J. Predicting Protein Function and Binding Profile via Matching of Local Evolutionary and Geometric Surface Patterns. Journal of Molecular Biology 387, 451–464 (2009).

21. Tseng, Y. Y., Chen, Z. J. & Li, W.-H. f POP: footprinting functional pockets of proteins by comparative spatial patterns. Nucleic Acids Research 38, D288–D295 (2010).

22. Xie, L., Xie, L. & Bourne, P. E. A unified statistical model to support local sequence order independent similarity searching for ligand-binding sites and its application to genome-based drug discovery. Bioinformatics 25, i305–i312 (2009).

23. Das, S., Krein, M. P. & Breneman, C. M. PESDserv: a server for high-throughput comparison of protein binding site surfaces. Bioinformatics 26, 1913–1914 (2010).

24. Standley, D. M., Yamashita, R., Kinjo, A. R., Toh, H. & Nakamura, H. SeSAW: balancing sequence and structural information in protein functional mapping. Bioinformatics 26, 1258–1259 (2010).

25. Moll, M., Bryant, D. H. & Kavraki, L. E. The LabelHash algorithm for substructure matching. BMC Bioinformatics 11, 555 (2010).

26. Weill, N. & Rognan, D. Alignment-Free Ultra-High-Throughput Comparison of Druggable Protein−Ligand Binding Sites. J. Chem. Inf. Model. 50, 123–135 (2010).

27. Desaphy, J., Azdimousa, K., Kellenberger, E. & Rognan, D. Comparison and Druggability Prediction of Protein–Ligand Binding Sites from Pharmacophore-Annotated Cavity Shapes. J. Chem. Inf. Model. 52, 2287–2299 (2012).

28. Wood, D. J., Vlieg, J. D., Wagener, M. & Ritschel, T. Pharmacophore Fingerprint-Based Approach to Binding Site Subpocket Similarity and Its Application to Bioisostere Replacement. J. Chem. Inf. Model. 52, 2031–2043 (2012).

29. Konc, J. & Janezic, D. ProBiS-2012: web server and web services for detection of structurally similar binding sites in proteins. Nucleic Acids Research 40, W214–W221 (2012).

30. Ito, J.-I., Tabei, Y., Shimizu, K., Tsuda, K. & Tomii, K. PoSSuM: a database of similar protein-ligand binding and putative pockets. Nucleic Acids Research 40, D541–D548 (2012).

31. Roy, A., Yang, J. & Zhang, Y. COFACTOR: an accurate comparative algorithm for structure-based protein function annotation. Nucleic Acids Research 40, W471–W477 (2012).

32. Nadzirin, N., Gardiner, E. J., Willett, P., Artymiuk, P. J. & Firdaus-Raih, M. SPRITE and ASSAM: web servers for side chain 3D-motif searching in protein structures. Nucleic Acids Research 40, W380–W386 (2012).

33. Lin, Y., Yoo, S. & Sanchez, R. SiteComp: a server for ligand binding site analysis in protein structures. Bioinformatics 28, 1172–1173 (2012).

34. Kurbatova, N., Chartier, M., Zylber, M. I. & Najmanovich, R. IsoCleft Finder – a web-based tool for the detection and analysis of protein binding-site geometric and chemical similarities. F1000Res 2, 117 (2013).

35. Kirshner, D. A., Nilmeier, J. P. & Lightstone, F. C. Catalytic site identification—a web server to identify catalytic site structural matches throughout PDB. Nucleic Acids Research 41, W256–W265 (2013).

36. Nadzirin, N., Willett, P., Artymiuk, P. J. & Firdaus-Raih, M. IMAAAGINE: a webserver for searching hypothetical 3D amino acid side chain arrangements in the Protein Data Bank. Nucleic Acids Research 41, W432–W440 (2013).

37. Gao, M. & Skolnick, J. APoc: large-scale identification of similar protein pockets. Bioinformatics 29, 597–604 (2013).

38. Desaphy, J., Raimbaud, E., Ducrot, P. & Rognan, D. Encoding Protein–Ligand Interaction Patterns in Fingerprints and Graphs. J. Chem. Inf. Model. 53, 623–637 (2013).

39. Caprari, S., Toti, D., Viet Hung, L., Di Stefano, M. & Polticelli, F. ASSIST: a fast versatile local structural comparison tool. Bioinformatics 30, 1022–1024 (2014).

40. Batista, J., Hawkins, P. C., Tolbert, R. & Geballe, M. T. SiteHopper - a unique tool for binding site comparison. J Cheminform 6, P57, 1758-2946-6-S1-P57 (2014).

41. Chartier, M., Adriansen, E. & Najmanovich, R. IsoMIF Finder: online detection of binding site molecular interaction field similarities. Bioinformatics 32, 621–623 (2016).

42. Chartier, M. & Najmanovich, R. Detection of Binding Site Molecular Interaction Field Similarities. J. Chem. Inf. Model. 55, 1600–1615 (2015).

43. Krotzky, T., Grunwald, C., Egerland, U. & Klebe, G. Large-Scale Mining for Similar Protein Binding Pockets: With RAPMAD Retrieval on the Fly Becomes Real. J. Chem. Inf. Model. 55, 165–179 (2015).

44. Lee, H. S. & Im, W. G-LoSA: An efficient computational tool for local structure-centric biological studies and drug design: G-LoSA. Protein Science 25, 865–876 (2016).

45. Núñez-Vivanco, G., Valdés-Jiménez, A., Besoaín, F. & Reyes-Parada, M. Geomfinder: a multi-feature identifier of similar three-dimensional protein patterns: a ligand-independent approach. Cheminform 8, 19 (2016).

46. Rey, J., Rasolohery, I., Tufféry, P., Guyon, F. & Moroy, G. PatchSearch: a web server for off-target protein identification. Nucleic Acids Research 47, W365–W372 (2019).

47. Ab Ghani, N. S., Ramlan, E. I. & Firdaus-Raih, M. Drug ReposER: a web server for predicting similar amino acid arrangements to known drug binding interfaces for potential drug repositioning. Nucleic Acids Research 47, W350–W356 (2019).

48. Simonovsky, M. & Meyers, J. DeeplyTough: Learning Structural Comparison of Protein Binding Sites. J. Chem. Inf. Model. 60, 2356–2366 (2020).

49. Breslin, P. A. S. An Evolutionary Perspective on Food and Human Taste. Current Biology 23, R409–R418 (2013).

50. Ho, H. K., et al. Functionally expressed bitter taste receptor TAS2R14 in human epidermal keratinocytes serves as a chemosensory receptor. Exp Dermatol 30, 216–225 (2021).

51. Shaw, L., et al. Personalized expression of bitter ‘taste’ receptors in human skin. PLoS ONE 13, e0205322 (2018).

52. Singh, N., Vrontakis, M., Parkinson, F. & Chelikani, P. Functional bitter taste receptors are expressed in brain cells. Biochemical and Biophysical Research Communications 406, 146–151 (2011).

53. Kyriazis, G. A., Soundarapandian, M. M. & Tyrberg, B. Sweet taste receptor signaling in beta cells mediates fructose-induced potentiation of glucose-stimulated insulin secretion. Proc. Natl. Acad. Sci. U.S.A. 109, (2012).

54. Kyriazis, G. A., Smith, K. R., Tyrberg, B., Hussain, T. & Pratley, R. E. Sweet Taste Receptors Regulate Basal Insulin Secretion and Contribute to Compensatory Insulin Hypersecretion During the Development of Diabetes in Male Mice. Endocrinology 155, 2112–2121 (2014).

55. Foster, S. R., et al. Expression, Regulation and Putative Nutrient-Sensing Function of Taste GPCRs in the Heart. PLoS ONE 8, e64579 (2013).

56. Foster, S. R., et al. Bitter taste receptor agonists elicit G[protein[dependent negative inotropy in the murine heart. FASEB j. 28, 4497–4508 (2014).

57. Deckmann, K., et al. Bitter triggers acetylcholine release from polymodal urethral chemosensory cells and bladder reflexes. Proc. Natl. Acad. Sci. U.S.A. 111, 8287–8292 (2014).

58. Deshpande, D. A., et al. Bitter taste receptors on airway smooth muscle bronchodilate by localized calcium signaling and reverse obstruction. Nature Medicine 16, 1299–1304 (2010).

59. Avau, B., et al. Targeting extra-oral bitter taste receptors modulates gastrointestinal motility with effects on satiation. Scientific Reports 5, 1–12 (2015).

60. Iwatsuki, K. & Uneyama, H. Sense of Taste in the Gastrointestinal Tract. J Pharmacol Sci 118, 123–128 (2012).

61. Raka, F., Farr, S., Kelly, J., Stoianov, A. & Adeli, K. Metabolic control via nutrient-sensing mechanisms: role of taste receptors and the gut-brain neuroendocrine axis. American Journal of Physiology-Endocrinology and Metabolism 317, E559–E572 (2019).

62. Carey, R. M. & Lee, R. J. Taste Receptors in Upper Airway Innate Immunity. Nutrients 11, 2017 (2019).

63. Behrens, M. & Lang, T. Extra-Oral Taste Receptors—Function, Disease, and Perspectives. Front. Nutr. 9, 881177 (2022).

64. Zhai, K., et al. Activation of bitter taste receptors (tas2rs) relaxes detrusor smooth muscle and suppresses overactive bladder symptoms. Oncotarget 7, 21156–21167 (2016).

65. Chandrashekar, J., et al. T2Rs Function as Bitter Taste Receptors. Cell 100, 703–711 (2000).

66. Sutkeviciute, I. & Vilardaga, J.-P. Structural insights into emergent signaling modes of G protein–coupled receptors. Journal of Biological Chemistry 295, 11626–11642 (2020).

67. Di Pizio, A. et al. Comparing Class A GPCRs to bitter taste receptors. in Biophysical Methods in Cell Biology vol. 132 401–427 (Elsevier Ltd, 2016).

68. Brockhoff, A., Behrens, M., Niv, M. Y. & Meyerhof, W. Structural requirements of bitter taste receptor activation. Proceedings of the National Academy of Sciences of the United States of America 107, 11110–11115 (2010).

69. Slack, J. P., et al. Modulation of Bitter Taste Perception by a Small Molecule hTAS2R Antagonist. Current Biology 20, 1104–1109 (2010).

70. Meyerhof, W., et al. The molecular receptive ranges of human TAS2R bitter taste receptors. Chemical Senses 35, 157–170 (2009).

71. Gowers, R. et al. MDAnalysis: A Python Package for the Rapid Analysis of Molecular Dynamics Simulations. in 98–105 (Austin, Texas, 2016). doi:10.25080/Majora-629e541a-00e.

72. Xu, W., et al. Structural basis for strychnine activation of human bitter taste receptor TAS2R46. Science 377, 1298–1304 (2022).

73. Born, S., Levit, A., Niv, M. Y., Meyerhof, W. & Behrens, M. The human bitter taste receptor TAS2R10 is tailored to accommodate numerous diverse ligands. Journal of Neuroscience 33, 201–213 (2013).

74. Jensen, A. A., Gharagozloo, P., Birdsall, N. J. M. & Zlotos, D. P. Pharmacological characterisation of strychnine and brucine analogues at glycine and α7 nicotinic acetylcholine receptors. European Journal of Pharmacology 539, 27–33 (2006).

75. DiPizio, A., et al. Rational design of agonists for bitter taste receptor TAS2R14: from modeling to bench and back. Cellular and Molecular Life Sciences 77, (2020).

76. Fierro, F., Giorgetti, A., Carloni, P., Meyerhof, W. & Alfonso-Prieto, M. Dual binding mode of “bitter sugars” to their human bitter taste receptor target. Scientific Reports 9, 1–16 (2019).

77. Nicoli, A., Dunkel, A., Giorgino, T., de Graaf, C. & Di Pizio, A. Classification Model for the Second Extracellular Loop of Class A GPCRs. J. Chem. Inf. Model. 62, 511–522 (2022).

78. Topin, J., et al. Functional molecular switches of mammalian G protein-coupled bitter-taste receptors. Cellular and Molecular Life Sciences 6, 2020.10.23.348706 (2021).

79. Kuipers, W., et al. Study of the interaction between aryloxypropanolamines and Asn386 in helix VII of the human 5-hydroxytryptamine1A receptor. Mol Pharmacol 51, 889–896 (1997).

80. Oksenberg, D., et al. A single amino-acid difference confers major pharmacological variation between human and rodent 5-HT1B receptors. Nature 360, 161–163 (1992).

81. Suryanarayana, S., Daunt, D. A., Von Zastrow, M. & Kobilka, B. K. A point mutation in the seventh hydrophobic domain of the alpha 2 adrenergic receptor increases its affinity for a family of beta receptor antagonists. J Biol Chem 266, 15488–15492 (1991).

82. Suryanarayana, S. & Kobilka, B. K. Amino acid substitutions at position 312 in the seventh hydrophobic segment of the beta 2-adrenergic receptor modify ligand-binding specificity. Mol Pharmacol 44, 111–114 (1993).

83. Erb, L., et al. Site-directed mutagenesis of P2U purinoceptors. Positively charged amino acids in transmembrane helices 6 and 7 affect agonist potency and specificity. J Biol Chem 270, 4185–4188 (1995).

84. Jiang, Q., et al. A Mutational Analysis of Residues Essential for Ligand Recognition at the Human P2Y_1_ Receptor. Mol Pharmacol 52, 499–507 (1997).

85. Kopin, A. S., McBride, E. W., Quinn, S. M., Kolakowski, L. F. & Beinborn, M. The role of the cholecystokinin-B/gastrin receptor transmembrane domains in determining affinity for subtype-selective ligands. J Biol Chem 270, 5019–5023 (1995).

86. Pronin, A. N. Identification of Ligands for Two Human Bitter T2R Receptors. Chemical Senses 29, 583–593 (2004).

87. Kuijpers, G. A. J., Vergara, L. A., Calvo, S. & Yadid, G. Inhibitory effect of strychnine on acetylcholine receptor activation in bovine adrenal medullary chromaffin cells. British J Pharmacology 113, 471–478 (1994).

88. García-Colunga, J. & Miledi, R. Modulation of nicotinic acetylcholine receptors by strychnine. Proc. Natl. Acad. Sci. U.S.A. 96, 4113–4118 (1999).

89. Zlotos, D. P., Mandour, Y. M. & Jensen, A. A. Strychnine and its mono- and dimeric analogues: a pharmaco-chemical perspective. Nat. Prod. Rep. 39, 1910–1937 (2022).

90. Abraham, M. J., et al. GROMACS: High performance molecular simulations through multi-level parallelism from laptops to supercomputers. SoftwareX 1–2, 19–25 (2015).

91. Rodrigues, J. P. G. L. M., Teixeira, J. M. C., Trellet, M. & Bonvin, A. M. J. J. pdb-tools: a swiss army knife for molecular structures. F1000Res 7, 1961 (2018).

92. Adasme, M. F. et al. PLIP 2021: expanding the scope of the protein–ligand interaction profiler to DNA and RNA. Nucleic Acids Research 49, W530–W534 (2021).

93. Spriggs, R. V., Artymiuk, P. J. & Willett, P. Searching for Patterns of Amino Acids in 3D Protein Structures. J. Chem. Inf. Comput. Sci. 43, 412–421 (2003).

94. Bron, C. & Kerbosch, J. Algorithm 457: finding all cliques of an undirected graph. Commun. ACM 16, 575–577 (1973).

95. Korb, O., Stützle, T. & Exner, T. E. PLANTS: Application of Ant Colony Optimization to Structure-Based Drug Design. in Ant Colony Optimization and Swarm Intelligence (eds. Dorigo, M. et al.) vol. 4150 247–258 (Springer Berlin Heidelberg, Berlin, Heidelberg, 2006).

96. Korb, O., Stützle, T. & Exner, T. E. An ant colony optimization approach to flexible protein– ligand docking. Swarm Intell 1, 115–134 (2007).

97. Çınaroğlu, S. S. & Timuçin, E. Comparative Assessment of Seven Docking Programs on a Nonredundant Metalloprotein Subset of the PDBbind Refined. J. Chem. Inf. Model. 59, 3846–3859 (2019).

98. Eldridge, M. D., Murray, C. W., Auton, T. R., Paolini, G. V. & Mee, R. P. Empirical scoring functions: I. The development of a fast empirical scoring function to estimate the binding affinity of ligands in receptor complexes. Journal of computer-aided molecular design 11, 425–45 (1997).

99. Sirci, F., et al. Virtual Fragment Screening: Discovery of Histamine H_3_ Receptor Ligands Using Ligand-Based and Protein-Based Molecular Fingerprints. J. Chem. Inf. Model. 52, 3308–3324 (2012).

100. Kooistra, A. J., Leurs, R., De Esch, I. J. P. & De Graaf, C. Structure-Based Prediction of G-Protein-Coupled Receptor Ligand Function: A β-Adrenoceptor Case Study. J. Chem. Inf. Model. 55, 1045–1061 (2015).

101. Bassani, D., Pavan, M., Sturlese, M. & Moro, S. Sodium or Not Sodium: Should Its Presence Affect the Accuracy of Pose Prediction in Docking GPCR Antagonists? Pharmaceuticals 15, 346 (2022).

102. Kooistra, A. J., et al. Function-specific virtual screening for GPCR ligands using a combined scoring method. Sci Rep 6, 28288 (2016).

103. De Graaf, C., et al. Crystal Structure-Based Virtual Screening for Fragment-like Ligands of the Human Histamine H_1_ Receptor. J. Med. Chem. 54, 8195–8206 (2011).

104. Huang, D. W., Sherman, B. T. & Lempicki, R. A. Systematic and integrative analysis of large gene lists using DAVID bioinformatics resources. Nat Protoc 4, 44–57 (2009).

105. Sherman, B. T. et al. DAVID: a web server for functional enrichment analysis and functional annotation of gene lists (2021 update). Nucleic Acids Research 50, W216–W221 (2022).

106. The Gene Ontology Consortium. The Gene Ontology project in 2008. Nucleic Acids Research 36, D440–D444 (2008).

107. Gillespie, M. et al. The reactome pathway knowledgebase 2022. Nucleic Acids Research 50, D687–D692 (2022).

108. Kanehisa, M. KEGG: Kyoto Encyclopedia of Genes and Genomes. Nucleic Acids Research 28, 27–30 (2000).

109. Kanehisa, M., Furumichi, M., Sato, Y., Kawashima, M. & Ishiguro-Watanabe, M. KEGG for taxonomy-based analysis of pathways and genomes. Nucleic Acids Research 51, D587–D592 (2023).

110. Ferreira, J. A. & Zwinderman, A. H. On the Benjamini–Hochberg method. The Annals of Statistics 34, 1827–1849 (2006).

111. NCBI Resource Coordinators et al. Database resources of the National Center for Biotechnology Information. Nucleic Acids Research 46, D8–D13 (2018).

112. Berman, H. M. The Protein Data Bank. Nucleic Acids Research 28, 235–242 (2000).

113. Jumper, J., et al. Highly accurate protein structure prediction with AlphaFold. Nature 596, 583–589 (2021).

114. Molecular Operating Environment (MOE), 2022.02 Chemical Computing Group ULC, 1010 Sherbooke St. West, Suite #910, Montreal, QC, Canada, H3A 2R7, 2023. (2022).

115. Jo, S., Kim, T., Iyer, V. G. & Im, W. CHARMM-GUI: A web-based graphical user interface for CHARMM. J. Comput. Chem. 29, 1859–1865 (2008).

116. Dickson, C. J., Walker, R. C. & Gould, I. R. Lipid21: Complex Lipid Membrane Simulations with AMBER. J. Chem. Theory Comput. 18, 1726–1736 (2022).

117. Tian, C., et al. ff19SB: Amino-Acid-Specific Protein Backbone Parameters Trained against Quantum Mechanics Energy Surfaces in Solution. J. Chem. Theory Comput. 16, 528–552 (2020).

118. Wang, J., Wolf, R. M., Caldwell, J. W., Kollman, P. A. & Case, D. A. Development and testing of a general amber force field. Journal of Computational Chemistry 25, 1157–1174 (2004).

119. Berendsen, H. J. C., Postma, J. P. M., van Gunsteren, W. F., DiNola, A. & Haak, J. R. Molecular dynamics with coupling to an external bath. The Journal of Chemical Physics 81, 3684–3690 (1984).

120. Nosé, S. A unified formulation of the constant temperature molecular dynamics methods. The Journal of Chemical Physics 81, 511–519 (1984).

121. Parrinello, M. & Rahman, A. Polymorphic transitions in single crystals: A new molecular dynamics method. Journal of Applied Physics 52, 7182–7190 (1981).

122. Hess, B., Bekker, H., Berendsen, H. J. C. & Fraaije, J. G. E. M. LINCS: A linear constraint solver for molecular simulations. Journal of Computational Chemistry 18, 1463–1472 (1997).

123. Humphrey, W., Dalke, A. & Schulten, K. VMD: Visual molecular dynamics. Journal of Molecular Graphics 14, 33–38 (1996).

