## Supplementary Information for "VirtuousPocketome: A Computational Tool for Screening Protein-ligand Complexes to Identify Similar Binding Sites"

[6] 7hc srl, Rome, Italy

*

† Lorenzo Pallante and Marco Cannariato contributed equally to this study.

### Supporting Information

**Results – Energy Minimisation and Conformational Dynamics**

Molecular dynamics simulations of the protein-ligand complex were performed according to the Methods section of the manuscript. In this section, we provided some details concerning the energy minimisation protocol and the convergence analysis of the conformational dynamics during simulations.

The minimization of the system converged after 2705 steps with a maximum force below 1000 kJ/(mol*nm) (Figure S1).

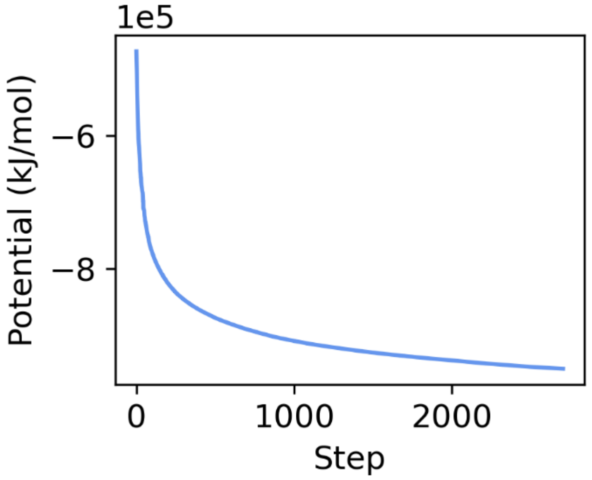


**Figure S1.** Potential energy of the simulated system during energy minimization. Energy converged after 2705 steps with a maximum force below 1000 kJ/(mol*nm).

To ensure the convergence of the MD simulations, we evaluated the RMSD and the cluster analysis as measures of simulation equilibrium for all the MD replicas. In detail, we calculated the RMSD of the protein backbone and the number of clusters during the last 50 ns using the linkage method with an RMSD cutoff of 0.15 nm. The RMSD trends reached a plateau (Figure S2) and only a single cluster was obtained during the last part of the simulation for all the replicas.

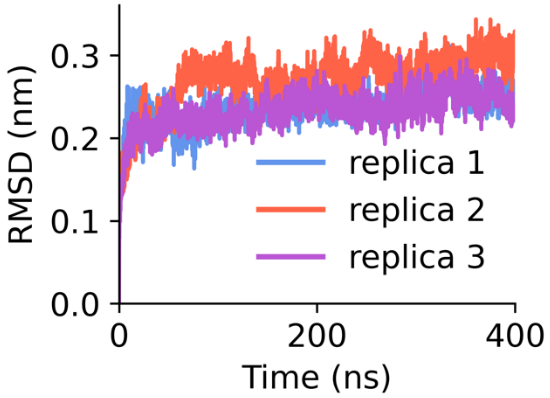


**Figure S2.** RMSD of the bitter taste receptor bound with strychnine for the three simulation replicas performed (colored in blue, red and violet, respectively).

According to these analyses, the last 50 ns of each simulation replica were considered as structural equilibrium and were concatenated to obtain a final 150 ns-long trajectory representing the ensemble of protein conformations.

#### Results – Motifs Creation

To characterise the protein-ligand interaction during the MD trajectory, we calculated the residue-specific probability interactions with strychnine. We used PLIP to calculate the frame-by-frame interactions during the concatenated trajectory (500 ns x 3 replicas = 1500 ns) analysing one frame every 200 ps. We then obtained the probability by dividing the count of each interaction by the total number of frames considered. Results are summarised in Table S1 and Figure S3.

**Table S1.** Interaction probabilities of protein residues with strychnine during the concatenated MD trajectory (500 ns x 3 replicas = 1500 ns). Residues TYR85, TRP88, and GLU265 demonstrated the highest values with interaction probability higher than 80% and are highlighted in light grey.

| Residue Name | Residue Number | Interaction Type | Probability |
|---|---|---|---|
| VAL | 61 | HI | 0.10 |
| LEU | 62 | HI | 0.37 |
| ASN | 65 | HB | 0.06 |
| ASN | 65 | HI | 0.35 |
| THR | 69 | HI | 0.16 |
| ARG | 81 | HI | 0.04 |
| ILE | 82 | HI | 0.01 |
| ALA | 84 | HI | 0.09 |
| TYR | 85 | HB | 0.07 |
| TYR | 85 | PI | 0.18 |
| TYR | 85 | HI | 0.82 |
| TRP | 88 | PI | 0.23 |
| TRP | 88 | HI | 0.93 |
| TYR | 173 | HB | 0.01 |
| ILE | 245 | HI | 0.01 |
| SER | 248 | HB | 0.23 |
| VAL | 249 | HI | 0.09 |
| PHE | 252 | HI | 0.37 |
| PHE | 261 | HI | 0.30 |

| | | | |
|---|---|---|---|
| GLU | 265 | HI | 0.07 |
| GLU | 265 | SB | 0.88 |
| ALA | 268 | HI | 0.11 |

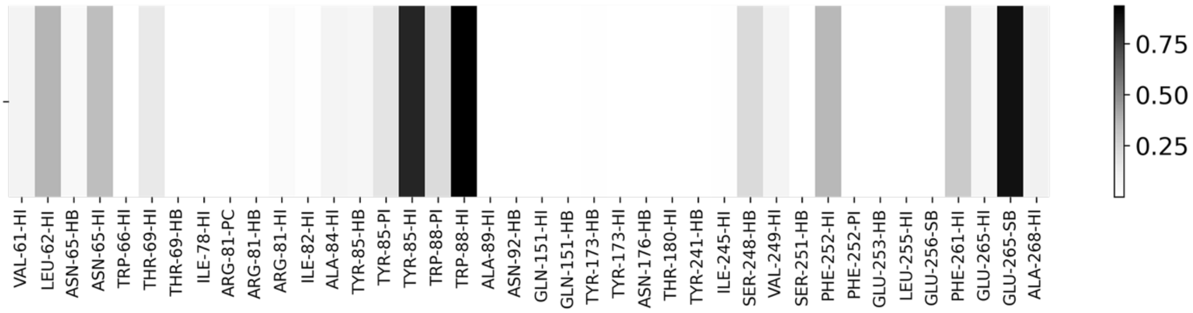


**Figure S3.** Interaction probabilities of protein residues with strychnine during the concatenated MD trajectory (500 ns x 3 replicas = 1500 ns). Probability values range from 0 to 1 and a grayscale colorbar was used, with light colours for low probability values and dark for high values. Residues TYR85, TRP88, and GLU265 demonstrated the highest values with interaction probability higher than 80% and are highlighted in dark grey.

#### Results - Similarity Search and Multi-step Filtering

The retrieved protein hits according to the docking score (DScore) after the multi-step filtering process are reported in Table S2.

**Table S2.** Protein hits according to the docking score (DScore) after the multi-step filtering process.

| PDB | Description | RMSD | SASA | DScore |
|---|---|---|---|---|
| **3a4s** | SUMO-CONJUGATING ENZYME UBC9 | 1.07 | 2.215 | -87.7711 |
| **3b7r** | LEUKOTRIENE A-4 HYDROLASE | 1.25 | 2.155 | -87.065 |
| **7bqz** | SPINDLIN-1 | 0.75 | 2.278 | -86.6839 |
| **2yg2** | APOLIPOPROTEIN M | 1.26 | 2.451 | -86.1774 |
| **3fhe** | LEUKOTRIENE A-4 HYDROLASE | 1.27 | 1.589 | -85.2467 |
| **2zfh** | CUTA | 1.36 | 1.936 | -85.0248 |
| **4dpr** | LEUKOTRIENE A-4 HYDROLASE | 1.31 | 2.073 | -85.0148 |
| **2r3r** | CELL DIVISION PROTEIN KINASE 2 | 1.13 | 2.306 | -84.9421 |
| **6o5h** | LEUKOTRIENE A-4 HYDROLASE | 1.3 | 2.348 | -84.9287 |
| **3fuk** | LEUKOTRIENE A-4 HYDROLASE | 1.26 | 1.86 | -84.7628 |
| **3fu5** | LEUKOTRIENE A-4 HYDROLASE | 1.29 | 2.97 | -84.591 |
| **7av2** | LEUKOTRIENE A-4 HYDROLASE | 1.26 | 1.684 | -84.3107 |
| **3fh7** | LEUKOTRIENE A-4 HYDROLASE | 1.22 | 2.443 | -84.2115 |
| **6r2u** | ZINC-ALPHA-2-GLYCOPROTEIN | 1.11 | 1.794 | -84.1411 |
| **6p5s** | HOMEODOMAIN-INTERACTING PROTEIN KINASE 2 | 1.16 | 2.097 | -84.1008 |
| **4r7l** | LEUKOTRIENE A-4 HYDROLASE | 1.25 | 1.298 | -84.0848 |
| **4ms6** | LEUKOTRIENE A-4 HYDROLASE | 1.34 | 1.489 | -83.9426 |
| **2h2u** | SOLUBLE CALCIUM-ACTIVATED NUCLEOTIDASE 1 | 1.02 | 2.114 | -83.8648 |
| **3chp** | LEUKOTRIENE A-4 HYDROLASE | 1.18 | 2.395 | -83.3851 |
| **7av1** | LEUKOTRIENE A-4 HYDROLASE | 1.29 | 1.685 | -83.3388 |
| **7kze** | LEUKOTRIENE A-4 HYDROLASE | 1.33 | 1.91 | -83.2739 |
| **5bpp** | LEUKOTRIENE A-4 HYDROLASE | 1.26 | 1.274 | -82.9787 |
| **3cho** | LEUKOTRIENE A-4 HYDROLASE | 1.24 | 1.414 | -82.8956 |
| **5ni4** | LEUKOTRIENE A-4 HYDROLASE | 1.26 | 2.004 | -82.8657 |
| **1hs6** | LEUKOTRIENE A-4 HYDROLASE | 1.26 | 2.677 | -82.8356 |
| **3ftv** | LEUKOTRIENE A-4 HYDROLASE | 1.24 | 1.533 | -82.6407 |
| **5ni6** | LEUKOTRIENE A-4 HYDROLASE | 1.23 | 2.295 | -82.4868 |
| **2zi2** | THROMBIN HEAVY CHAIN | 1.16 | 2.086 | -82.4733 |
| **7auz** | LEUKOTRIENE A-4 HYDROLASE | 1.28 | 1.45 | -82.4511 |
| **6end** | LEUKOTRIENE A-4 HYDROLASE | 1.25 | 2.812 | -82.2884 |
| **3ftu** | LEUKOTRIENE A-4 HYDROLASE | 1.25 | 2.925 | -82.141 |
| **3ful** | LEUKOTRIENE A-4 HYDROLASE | 1.31 | 1.81 | -82.1373 |
| **7qg0** | NAD(+) HYDROLASE SARM1 | 1.39 | 3.449 | -81.9657 |
| **4rsy** | LEUKOTRIENE A-4 HYDROLASE | 1.26 | 1.774 | -81.922 |
| **7n3n** | SOLUTE CARRIER FAMILY 12 MEMBER 2 | 1.26 | 2.091 | -81.8164 |
| **2r24** | ALDOSE REDUCTASE | 1.37 | 1.567 | -81.7647 |
| **1gw6** | LEUKOTRIENE A-4 HYDROLASE | 1.24 | 2.231 | -81.7538 |
| **6enb** | LEUKOTRIENE A-4 HYDROLASE | 1.28 | 1.961 | -81.7255 |
| **5kir** | PROSTAGLANDIN G/H SYNTHASE 2 | 1.43 | 2.149 | -81.1718 |

| | | | | |
|---|---|---|---|---|
| **4l2l** | LEUKOTRIENE A-4 HYDROLASE | 1.3 | 1.311 | -81.1157 |
| **3eq6** | ACYL-COENZYME A SYNTHETASE ACSM2A | 1.27 | 2.577 | -80.8085 |
| **3fu3** | LEUKOTRIENE A-4 HYDROLASE | 1.29 | 3.235 | -80.7407 |
| **2vqo** | HISTONE DEACETYLASE 4 | 1.06 | 2.148 | -80.5489 |
| **6p8z** | GTPASE KRAS | 1.34 | 2.009 | -80.5234 |
| **3fty** | LEUKOTRIENE A-4 HYDROLASE | 1.25 | 2.088 | -80.4234 |
| **3chs** | LEUKOTRIENE A-4 HYDROLASE | 1.21 | 1.943 | -80.4112 |
| **6w9v** | TCR-BETA CHAIN, MAJOR HISTOCOMPATIBILITY COMPLEX CLASS I-RELATED GENE | 1.37 | 2.865 | -80.3695 |
| **2r3o** | CELL DIVISION PROTEIN KINASE 2 | 1.28 | 1.456 | -79.8837 |
| **6h0g** | PROTEIN CEREBLON, DNA DAMAGE-BINDING PROTEIN 1,DNA DAMAGE-BINDING PROTEIN 1, | 1.45 | 2.141 | -79.6843 |
| **7ure** | ISOFORM 2 OF PROTEIN-SERINE O-PALMITOLEOYLTRANSFERASE | 1.22 | 1.97 | -79.589 |
| **4kfz** | ANTI-LMO2 VH | 1.49 | 2.086 | -79.5841 |
| **3mph** | AMILORIDE-SENSITIVE AMINE OXIDASE | 1.48 | 1.988 | -79.4227 |
| **6u3p** | ACETYLCHOLINESTERASE | 1.5 | 1.988 | -79.3888 |
| **5fe7** | HISTONE ACETYLTRANSFERASE KAT2B | 1.42 | 1.618 | -78.6233 |
| **6nr8** | T-COMPLEX PROTEIN 1 SUBUNIT GAMMA, T-COMPLEX PROTEIN 1 SUBUNIT THETA | 1.47 | 1.547 | -78.5151 |
| **6m5o** | SERINE HYDROXYMETHYLTRANSFERASE, MITOCHONDRIAL | 1.37 | 1.652 | -78.2072 |
| **5mw3** | HISTONE-LYSINE N-METHYLTRANSFERASE, H3 LYSINE-79 SPECIFIC | 1.36 | 1.407 | -78.0836 |
| **3d49** | THROMBIN HEAVY CHAIN | 1.16 | 1.609 | -78.0821 |
| **1msv** | S-ADENOSYLMETHIONINE DECARBOXYLASE PROENZYME | 1.44 | 1.902 | -77.9728 |
| **5fe1** | HISTONE ACETYLTRANSFERASE KAT2B | 1.4 | 1.399 | -77.9383 |
| **3qrt** | CYCLIN-DEPENDENT KINASE 2 | 1.45 | 2.434 | -77.9 |
| **3qvv** | SULFOTRANSFERASE 1A1 | 1.45 | 1.562 | -77.8891 |
| **7zjp** | TRANSCRIPTIONAL ENHANCER FACTOR TEF-1 | 1.46 | 2.805 | -77.8122 |
| **4umo** | CALMODULIN, POTASSIUM VOLTAGE-GATED CHANNEL SUBFAMILY KQT MEMBER 1 | 1.23 | 1.668 | -77.6608 |
| **7e9n** | HEAVY CHAIN OF 35B5 FAB | 1.15 | 2.144 | -77.4416 |
| **6f5t** | LYSINE-SPECIFIC DEMETHYLASE 4D | 1.45 | 1.785 | -77.2974 |
| **5m7t** | PROTEIN O-GLCNACASE | 1.37 | 2.185 | -77.279 |
| **1sqm** | LEUKOTRIENE A-4 HYDROLASE | 1.29 | 1.249 | -77.1773 |
| **8hkw** | PEPTIDE FROM TP53-BINDING PROTEIN 1, IMPORTIN SUBUNIT ALPHA-3 | 1.47 | 1.519 | -76.9298 |
| **3lnz** | E3 UBIQUITIN-PROTEIN LIGASE MDM2 | 1.45 | 2.599 | -76.7359 |
| **4ek3** | CYCLIN-DEPENDENT KINASE 2 | 1.22 | 1.817 | -76.706 |
| **3ibd** | CYTOCHROME P450 2B6 | 1.43 | 1.746 | -76.7021 |
| **2l12** | CHROMOBOX HOMOLOG 7 | 1.25 | 3.816 | -76.2799 |
| **3rvh** | LYSINE-SPECIFIC DEMETHYLASE 4A | 1.29 | 1.531 | -76.2637 |
| **6qnx** | COHESIN SUBUNIT SA-2, TRANSCRIPTIONAL REPRESSOR CTCF | 1.18 | 2.616 | -76.2505 |
| **6hg4** | INTERLEUKIN-17 RECEPTOR C | 1.48 | 1.43 | -75.8588 |
| **4nst** | CYCLIN-DEPENDENT KINASE 12, CYCLIN-K | 1.46 | 2.276 | -75.8423 |
| **4x0u** | ALPHA-AMINOADIPIC SEMIALDEHYDE DEHYDROGENASE | 1.42 | 2.124 | -75.8376 |
| **7bmk** | SERINE/THREONINE-PROTEIN KINASE/ENDORIBONUCLEASE IRE1 | 1.38 | 3.453 | -75.8372 |
| **4bbm** | CYCLIN-DEPENDENT KINASE-LIKE 2 | 1.31 | 1.714 | -75.8371 |
| **2e8d** | BETA-2-MICROGLOBULIN | 1.49 | 2.138 | -75.811 |
| **4xx1** | FAB1 HEAVY CHAIN | 1.32 | 1.994 | -75.8052 |
| **6yaf** | AP-2 COMPLEX SUBUNIT BETA | 1.32 | 1.496 | -75.7476 |
| **6qpl** | SPINDLIN-1 | 1.45 | 2.241 | -75.724 |
| **4mzg** | SPINDLIN-1 | 1.41 | 2.286 | -75.6688 |
| **1gz8** | CELL DIVISION PROTEIN KINASE 2 | 1.19 | 3.287 | -75.6682 |
| **6j0l** | BUTYROPHILIN SUBFAMILY 3 MEMBER A3 | 1.19 | 1.654 | -75.6497 |
| **6yah** | AP-2 COMPLEX SUBUNIT BETA | 1.32 | 1.466 | -75.5767 |
| **6mid** | MONOCLONAL ANTIBODY ZIKV-195 HEAVY CHAIN | 1.48 | 1.145 | -75.4824 |
| **6tel** | HISTONE-LYSINE N-METHYLTRANSFERASE, H3 LYSINE-79 SPECIFIC | 1.33 | 2.339 | -75.2809 |
| **7rhl** | CGMP-GATED CATION CHANNEL ALPHA-1 | 1.15 | 2.253 | -75.1955 |
| **7zyf** | LEUCYL-CYSTINYL AMINOPEPTIDASE, PREGNANCY SERUM FORM | 0.94 | 2.367 | -75.148 |
| **3u84** | MENIN | 1.22 | 2.05 | -75 |
| **7qne** | GABA(A) RECEPTOR SUBUNIT GAMMA-2, GAMMA-AMINOBUTYRIC ACID RECEPTOR SUBUNIT BETA-3 | 1.41 | 1.498 | -74.9855 |
| **6xk9** | PROTEIN CEREBLON, DNA DAMAGE-BINDING PROTEIN 1 | 1.38 | 1.429 | -74.8752 |
| **6vum** | STEROL O-ACYLTRANSFERASE 1 | 1.48 | 2.684 | -74.8556 |
| **7p9t** | 5'-NUCLEOTIDASE | 1.45 | 2.011 | -74.8448 |
| **8csq** | 28S RIBOSOMAL PROTEIN S29, MITOCHONDRIAL | 1.29 | 1.622 | -74.8231 |
| **1cly** | IGG FAB (HUMAN IGG1, KAPPA) | 1.23 | 1.784 | -74.6885 |
| **4u9v** | N-ALPHA-ACETYLTRANSFERASE 40 | 1.31 | 2.191 | -74.6182 |
| **4ljp** | E3 UBIQUITIN-PROTEIN LIGASE RNF31 | 1.46 | 2.269 | -74.4844 |
| **2r3h** | CELL DIVISION PROTEIN KINASE 2 | 1 | 2.291 | -74.3763 |
| **7byi** | SERINE HYDROXYMETHYLTRANSFERASE, MITOCHONDRIAL | 1.4 | 1.907 | -74.2523 |
| **7qoo** | CENTROMERE PROTEIN H, CENTROMERE PROTEIN I | 0.72 | 2.216 | -74.234 |
| **7ocb** | SPINDLIN-1 | 1.44 | 3.322 | -74.153 |
| **6v8c** | ORNITHINE AMINOTRANSFERASE, MITOCHONDRIAL | 1.34 | 1.647 | -73.9414 |
| **6in3** | HISTONE-LYSINE N-METHYLTRANSFERASE, H3 LYSINE-79 SPECIFIC | 1.33 | 3.301 | -73.8186 |

| | | | | |
|---|---|---|---|---|
| **6wqz** | AUTOPHAGY-RELATED PROTEIN 9A | 1.22 | 2.699 | -73.707 |
| **7r5s** | CENTROMERE PROTEIN H, CENTROMERE PROTEIN I | 0.96 | 3.945 | -73.6423 |
| **7ttn** | TUBULIN BETA CHAIN, T-COMPLEX PROTEIN 1 SUBUNIT GAMMA | 1.34 | 1.756 | -73.5768 |
| **1grh** | GLUTATHIONE REDUCTASE | 1.46 | 2.023 | -73.4928 |
| **1grg** | GLUTATHIONE REDUCTASE | 1.46 | 1.613 | -73.4914 |
| **3vv0** | HISTONE-LYSINE N-METHYLTRANSFERASE SETD7 | 1.19 | 1.978 | -73.342 |
| **5wbs** | FRIZZLED-7,INHIBITOR PEPTIDE FZ7-21 | 1.09 | 2.043 | -73.3394 |
| **3grs** | GLUTATHIONE REDUCTASE | 1.46 | 1.35 | -73.3385 |
| **7m63** | INDOLEAMINE 2,3-DIOXYGENASE 1 | 1.37 | 1.501 | -73.2505 |
| **4pvf** | SERINE HYDROXYMETHYLTRANSFERASE, MITOCHONDRIAL | 1.37 | 2.059 | -73.2492 |
| **8dwt** | SPECKLE-TYPE POZ PROTEIN | 1.33 | 2.068 | -73.2317 |
| **7omn** | JD1-1 VH DOMAIN | 1.35 | 1.915 | -73.2243 |
| **7q29** | ANGIOTENSIN-CONVERTING ENZYME | 1.5 | 2.346 | -73.2221 |
| **6bly** | CLEAVAGE AND POLYADENYLATION SPECIFICITY FACTOR SUBUNIT 1 | 1.38 | 1.682 | -73.1401 |
| **6ba4** | HISTONE ACETYLTRANSFERASE KAT8 | 1.4 | 2.132 | -73.089 |
| **5fpb** | LYSINE-SPECIFIC DEMETHYLASE 4D | 1.32 | 1.492 | -72.8148 |
| **1grf** | GLUTATHIONE REDUCTASE | 1.45 | 2.048 | -72.7912 |
| **2c8y** | THROMBIN HEAVY CHAIN | 1.17 | 1.538 | -72.7212 |
| **6mbl** | HISTONE-LYSINE N-METHYLTRANSFERASE SETD3 | 1.04 | 2.258 | -72.679 |
| **2vtm** | CELL DIVISION PROTEIN KINASE 2 | 1.44 | 1.645 | -72.5833 |
| **7o7l** | ALPHA-2-MACROGLOBULIN | 1.33 | 1.796 | -72.5426 |
| **1eak** | 72 KDA TYPE IV COLLAGENASE | 1.43 | 1.489 | -72.4855 |
| **5ja7** | CATHEPSIN K | 1.09 | 1.968 | -72.4568 |
| **5z9w** | EBOLAVIRUS NUCLEOPROTEIN (RESIDUES 19-406) | 0.83 | 1.86 | -72.4313 |
| **7lk0** | ORNITHINE AMINOTRANSFERASE, MITOCHONDRIAL | 1.35 | 2.123 | -72.3516 |
| **7fcp** | P5-22 ANTIBODY FAB FRAGMENT HEAVY CHAIN | 1.5 | 1.196 | -72.3062 |
| **6w3c** | SERINE/THREONINE-PROTEIN KINASE/ENDORIBONUCLEASE IRE1 | 1.34 | 1.72 | -72.2763 |
| **5bnj** | CYCLIN-DEPENDENT KINASE 8 | 1.32 | 1.454 | -72.2569 |
| **6xdb** | SERINE/THREONINE-PROTEIN KINASE/ENDORIBONUCLEASE IRE1 | 1.42 | 3.229 | -72.2502 |
| **7uvr** | ATP-DEPENDENT CLP PROTEASE PROTEOLYTIC SUBUNIT, | 1.24 | 1.621 | -72.2437 |
| **6i8b** | SPINDLIN-1 | 1.44 | 2.006 | -72.077 |
| **7qdr** | WD REPEAT-CONTAINING PROTEIN 61 | 1.47 | 2.207 | -71.9206 |
| **7nvl** | T-COMPLEX PROTEIN 1 SUBUNIT ETA | 1.25 | 1.451 | -71.8865 |
| **5jq8** | CYCLIN-DEPENDENT KINASE 2 | 1.24 | 1.522 | -71.813 |
| **7a3g** | DIPEPTIDYL PEPTIDASE 8 | 1.47 | 2.615 | -71.8083 |
| **5fe6** | HISTONE ACETYLTRANSFERASE KAT2B | 1.35 | 2.409 | -71.7955 |
| **5fbh** | EXTRACELLULAR CALCIUM-SENSING RECEPTOR | 1.34 | 1.521 | -71.7642 |
| **5fe2** | HISTONE ACETYLTRANSFERASE KAT2B | 1.34 | 1.605 | -71.7247 |
| **5mv7** | UNCONVENTIONAL MYOSIN-VIIB | 1.29 | 2.397 | -71.6435 |
| **5fy8** | LYSINE-SPECIFIC DEMETHYLASE 4A | 1.11 | 2.043 | -71.6353 |
| **6h4q** | LYSINE-SPECIFIC DEMETHYLASE 4A | 1.25 | 3.312 | -71.611 |
| **4udw** | THROMBIN HEAVY CHAIN | 1.23 | 2.865 | -71.5643 |
| **5y3r** | DNA-DEPENDENT PROTEIN KINASE CATALYTIC SUBUNIT | 1.24 | 2.279 | -71.5376 |
| **2ypt** | CAAX PRENYL PROTEASE 1 HOMOLOG | 1.31 | 2.139 | -71.5061 |
| **5ka8** | TYROSINE-PROTEIN PHOSPHATASE NON-RECEPTOR TYPE 1 | 0.99 | 1.969 | -71.4398 |
| **2rhy** | LETHAL(3)MALIGNANT BRAIN TUMOR-LIKE PROTEIN | 1.32 | 1.6 | -71.4055 |
| **8hik** | ANTI-BRIL FAB LIGHT CHAIN, ANTI-BRIL FAB HEAVY CHAIN | 1.25 | 2.178 | -71.4044 |
| **5uwf** | CAMP AND CAMP-INHIBITED CGMP 3',5'-CYCLIC PHOSPHODIESTERASE | 1.38 | 2.096 | -71.303 |
| **8e3i** | CLEAVAGE AND POLYADENYLATION SPECIFICITY FACTOR SUBUNIT 1 | 1.38 | 1.756 | -71.2577 |
| **7sck** | EXOSTOSIN-2 | 1.11 | 2.409 | -71.1007 |
| **2wjy** | REGULATOR OF NONSENSE TRANSCRIPTS 1 | 1.37 | 1.534 | -71.0623 |
| **4p4h** | MITOCHONDRIAL ANTIVIRAL-SIGNALING PROTEIN, PROBABLE ATP-DEPENDENT RNA HELICASE DDX58 | 1.24 | 1.584 | -71.0279 |
| **7ni5** | SERINE-PROTEIN KINASE ATM | 1.25 | 2.787 | -71.0278 |
| **5hnb** | CYCLIN-DEPENDENT KINASE 8 | 1.29 | 1.458 | -71.0242 |
| **5xez** | ANTIBODY, MAB1, HEAVY CHAIN | 1.43 | 1.533 | -71.014 |
| **7nd4** | COVOX-88 FAB HEAVY CHAIN | 1.47 | 2.694 | -70.9952 |
| **4ifb** | BILE SALT SULFOTRANSFERASE | 1.42 | 1.855 | -70.8993 |
| **7d0p** | HISTONE ACETYLTRANSFERASE KAT7 | 1.48 | 1.494 | -70.899 |
| **7nvn** | T-COMPLEX PROTEIN 1 SUBUNIT ETA | 1.22 | 1.425 | -70.8572 |
| **6r7n** | COP9 SIGNALOSOME COMPLEX SUBUNIT 2, CULLIN-2 | 1.21 | 2.267 | -70.8549 |
| **4tw0** | SCAVENGER RECEPTOR CLASS B MEMBER 2 | 1.42 | 2.458 | -70.8325 |
| **5q8h** | DCLRE1A | 1.46 | 1.94 | -70.8299 |
| **3epa** | S-ADENOSYLMETHIONINE DECARBOXYLASE BETA CHAIN, S-ADENOSYLMETHIONINE DECARBOXYLASE ALPHA CHAIN | 1.44 | 2.078 | -70.8049 |
| **1y1j** | C-ALPHA-FORMYGLYCINE-GENERATING ENZYME | 1.28 | 1.745 | -70.7845 |
| **6u8w** | DNA (CYTOSINE-5)-METHYLTRANSFERASE 3B, DNA (CYTOSINE-5)-METHYLTRANSFERASE 3-LIKE | 1.46 | 1.576 | -70.7426 |
| **7o7o** | ALPHA-2-MACROGLOBULIN | 1.23 | 1.581 | -70.7403 |
| **6rnq** | GEM-ASSOCIATED PROTEIN 5 | 0.84 | 1.908 | -70.736 |
| **6h4u** | LYSINE-SPECIFIC DEMETHYLASE 4A | 1.29 | 1.781 | -70.6779 |
| **4ayv** | THROMBIN HEAVY CHAIN | 1.29 | 1.922 | -70.6612 |
| **7ted** | ORNITHINE AMINOTRANSFERASE, MITOCHONDRIAL | 1.32 | 3.313 | -70.6456 |

| | | | | |
|---|---|---|---|---|
| **3zmz** | LYSINE-SPECIFIC HISTONE DEMETHYLASE 1A, REST COREPRESSOR 1 | 1.42 | 1.997 | -70.5755 |
| **4aw6** | CAAX PRENYL PROTEASE 1 HOMOLOG | 1.32 | 2.358 | -70.5584 |
| **8b6l** | TRANSMEMBRANE PROTEIN 258, DOLICHYL-DIPHOSPHOOLIGOSACCHARIDE--PROTEIN | 1.32 | 1.701 | -70.5459 |
| **5lsp** | 107_A07 FAB HEAVY CHAIN, 107_A07 FAB LIGHT CHAIN | 1.26 | 1.753 | -70.5312 |
| **5q8q** | DCLRE1A | 1.49 | 3.285 | -70.493 |
| **2xiq** | METHYLMALONYL-COA MUTASE, MITOCHONDRIAL | 1.45 | 1.836 | -70.4676 |
| **4msn** | CAMP AND CAMP-INHIBITED CGMP 3',5'-CYCLIC PHOSPHODIESTERASE | 1.44 | 2.813 | -70.4675 |
| **6s8l** | TUBULIN BETA-3 CHAIN | 1.12 | 1.729 | -70.4406 |
| **4ek9** | HISTONE-LYSINE N-METHYLTRANSFERASE, H3 LYSINE-79 SPECIFIC | 1.25 | 1.738 | -70.4255 |
| **7sid** | SERINE-PROTEIN KINASE ATM | 1.37 | 2.352 | -70.4108 |
| **7k8s** | C002 FAB HEAVY CHAIN, C002 FAB LIGHT CHAIN | 1.29 | 1.609 | -70.3647 |
| **5luq** | DNA-DEPENDENT PROTEIN KINASE CATALYTIC SUBUNIT,DNA- | 1.05 | 1.73 | -70.3546 |
| **7xur** | SNRNA-ACTIVATING PROTEIN COMPLEX SUBUNIT 3 | 1.4 | 1.978 | -70.3087 |
| **7ni6** | SERINE-PROTEIN KINASE ATM | 1.3 | 2.067 | -70.3043 |
| **7m30** | 1-32 FAB HEAVY CHAIN | 1.44 | 1.662 | -70.2799 |
| **5ffg** | INTEGRIN BETA-6, INTEGRIN ALPHA-V | 1.31 | 2.065 | -70.2695 |
| **6dv5** | HEAT SHOCK PROTEIN BETA-1 | 1.31 | 1.541 | -70.2648 |
| **7ye9** | LIGHT CHAIN OF R1-32 FAB | 1.18 | 2.219 | -70.2502 |
| **6d01** | 1210 ANTIBODY, LIGHT CHAIN, 1210 ANTIBODY, HEAVY CHAIN | 1.24 | 1.737 | -70.2299 |
| **7q12** | GLYCOGEN [STARCH] SYNTHASE, MUSCLE | 1.44 | 3.126 | -70.1953 |
| **6ugq** | CARBONIC ANHYDRASE IX-MIMIC | 1.24 | 1.4 | -70.1773 |
| **8aql** | SERINE HYDROXYMETHYLTRANSFERASE, MITOCHONDRIAL | 1.32 | 1.257 | -70.1732 |
| **7m7d** | INDOLEAMINE 2,3-DIOXYGENASE 1 | 1.34 | 1.995 | -70.1541 |
| **3apw** | ALPHA-1-ACID GLYCOPROTEIN 2 | 1.43 | 2.657 | -70.1114 |
| **4ekz** | PROTEIN DISULFIDE-ISOMERASE | 1.18 | 2.471 | -70.0843 |
| **1t2a** | GDP-MANNOSE 4,6 DEHYDRATASE | 1.47 | 1.527 | -70.0784 |
| **3b9f** | PROTHROMBIN | 1.1 | 2.144 | -70.0644 |
| **5q4w** | DCLRE1A | 1.47 | 1.934 | -70.0288 |
| **2hi8** | SULFATASE-MODIFYING FACTOR 1 | 1.29 | 1.953 | -70.0168 |
| **6vkg** | HUMAN CARBONIC ANHYDRASE IX MIMIC | 1.34 | 1.914 | -70.0096 |
| **3vxp** | HLA CLASS I HISTOCOMPATIBILITY ANTIGEN, A-24 ALPHA CHAIN | 1.49 | 1.638 | -69.9877 |
| **6v63** | ACTIN-HISTIDINE N-METHYLTRANSFERASE | 0.96 | 3.151 | -69.9484 |
| **7nvm** | T-COMPLEX PROTEIN 1 SUBUNIT ETA | 1.25 | 1.497 | -69.9173 |
| **2xij** | METHYLMALONYL-COA MUTASE, MITOCHONDRIAL | 1.46 | 2.278 | -69.8883 |
| **7zsc** | PROLYL 4-HYDROXYLASE SUBUNIT ALPHA-2, PROTEIN DISULFIDE-ISOMERASE | 1.21 | 1.972 | -69.7845 |
| **8e3q** | CLEAVAGE AND POLYADENYLATION SPECIFICITY FACTOR SUBUNIT 1 | 1.45 | 1.685 | -69.7186 |
| **1h4r** | MERLIN | 1.17 | 1.743 | -69.6992 |
| **7czx** | IG C168_LIGHT_IGKV4-1_IGKJ4,UNCHARACTERIZED PROTEIN | 1.49 | 1.843 | -69.6461 |
| **6bnb** | PROTEIN CEREBLON, DNA DAMAGE-BINDING PROTEIN 1 | 1.45 | 1.865 | -69.6368 |
| **6oht** | 3-BETA-HYDROXYSTEROID-DELTA(8),DELTA(7)-ISOMERASE | 1 | 1.92 | -69.5843 |
| **2ckj** | XANTHINE OXIDOREDUCTASE | 1 | 2.287 | -69.5344 |
| **6mie** | POTASSIUM VOLTAGE-GATED CHANNEL SUBFAMILY KQT MEMBER 1 | 1.34 | 2.218 | -69.5254 |
| **7fem** | ANGIOTENSIN-CONVERTING ENZYME 2 | 1.27 | 1.828 | -69.5141 |
| **6cqd** | MITOGEN-ACTIVATED PROTEIN KINASE KINASE KINASE KINASE 1 | 1.08 | 2.839 | -69.4974 |
| **2aii** | SULFATASE MODIFYING FACTOR 1 | 1.29 | 2.389 | -69.4813 |
| **2qrv** | DNA (CYTOSINE-5)-METHYLTRANSFERASE 3A, DNA (CYTOSINE-5)-METHYLTRANSFERASE 3-LIKE | 1.43 | 2.655 | -69.3916 |
| **7v88** | ANGIOTENSIN-CONVERTING ENZYME 2,ANGIOTENSIN-CONVERTING | 1.09 | 2.18 | -69.3707 |
| **4liq** | MACROPHAGE COLONY-STIMULATING FACTOR 1 RECEPTOR | 1.46 | 1.47 | -69.3014 |
| **6cxv** | INDOLEAMINE 2,3-DIOXYGENASE 1 | 1.44 | 2.39 | -69.2881 |
| **7rew** | ANTI-CYNO INTERLEUKIN 13 FAB HEAVY CHAIN | 1.44 | 1.912 | -69.2592 |
| **7q3n** | UROMODULIN | 1.32 | 3.582 | -69.2552 |
| **5vk0** | E3 UBIQUITIN-PROTEIN LIGASE MDM2 | 1.29 | 2.35 | -69.2364 |
| **2rhu** | LETHAL(3)MALIGNANT BRAIN TUMOR-LIKE PROTEIN | 1.31 | 3.016 | -69.2313 |
| **4x7t** | OMALIZUMAB-FAB HEAVY CHAIN | 1.48 | 1.304 | -69.226 |
| **6bb2** | L-LACTATE DEHYDROGENASE A CHAIN | 1.04 | 2.177 | -69.2087 |
| **5j13** | ANTI-TSLP FAB-FRAGMENT, HEAVY CHAIN | 0.93 | 1.614 | -69.1306 |
| **5a7p** | LYSINE-SPECIFIC DEMETHYLASE 4A | 1.11 | 2.043 | -69.0758 |
| **5l3c** | LYSINE-SPECIFIC HISTONE DEMETHYLASE 1A, REST COREPRESSOR 1 | 1.38 | 1.651 | -69.0717 |
| **8d7w** | PROTEIN CEREBLON, DNA DAMAGE-BINDING PROTEIN 1 | 1.47 | 1.444 | -69.0315 |
| **1igr** | INSULIN-LIKE GROWTH FACTOR RECEPTOR 1 | 1.5 | 1.697 | -68.9379 |
| **3tiy** | CYCLIN-DEPENDENT KINASE 2 | 1.15 | 1.536 | -68.9332 |
| **6v01** | POTASSIUM VOLTAGE-GATED CHANNEL SUBFAMILY KQT MEMBER 1 | 1.46 | 2.354 | -68.9038 |
| **3fz1** | CELL DIVISION PROTEIN KINASE 2 | 1.12 | 2.286 | -68.892 |
| **6ohs** | PHOSPHOLIPASE D2 | 1.44 | 2.056 | -68.8836 |
| **3m17** | IGG RECEPTOR FCRN LARGE SUBUNIT P51 | 1.44 | 2.255 | -68.8318 |
| **5fwe** | LYSINE-SPECIFIC DEMETHYLASE 4A | 1.34 | 1.823 | -68.8306 |
| **4u7p** | DNA (CYTOSINE-5)-METHYLTRANSFERASE 3A, DNA (CYTOSINE-5)-METHYLTRANSFERASE 3-LIKE | 1.45 | 2.049 | -68.8292 |
| **7wi0** | XMA01 HEAVY CHAIN VARIABLE DOMAIN | 1.44 | 2.458 | -68.8238 |
| **6b8z** | TYROSINE-PROTEIN PHOSPHATASE NON-RECEPTOR TYPE 1 | 0.89 | 2.064 | -68.8222 |

| | | | | |
|---|---|---|---|---|
| **2f27** | SIALIDASE 2 | 1.46 | 2.002 | -68.7455 |
| **3lfs** | CELL DIVISION PROTEIN KINASE 2 | 1.15 | 2.086 | -68.699 |
| **4btj** | TAU-TUBULIN KINASE 1 | 1.15 | 1.437 | -68.6537 |
| **7ul3** | HISTAMINE H2 RECEPTOR | 1.43 | 1.995 | -68.6327 |
| **4j9e** | TYROSINE-PROTEIN KINASE ABL1, P17 | 1.44 | 2.116 | -68.6259 |
| **5lvr** | HISTONE ACETYLTRANSFERASE KAT2B | 1.42 | 1.542 | -68.625 |
| **3k3o** | PHD FINGER PROTEIN 8 | 1.08 | 2.144 | -68.6182 |
| **7frf** | TYROSINE-PROTEIN PHOSPHATASE NON-RECEPTOR TYPE 1 | 1.19 | 2.226 | -68.6141 |
| **4az2** | THROMBIN HEAVY CHAIN | 1.18 | 2.583 | -68.5636 |
| **3smt** | HISTONE-LYSINE N-METHYLTRANSFERASE SETD3 | 0.92 | 2.244 | -68.5505 |

#### Results - Functional Enrichment and Signalling Pathway Analyses

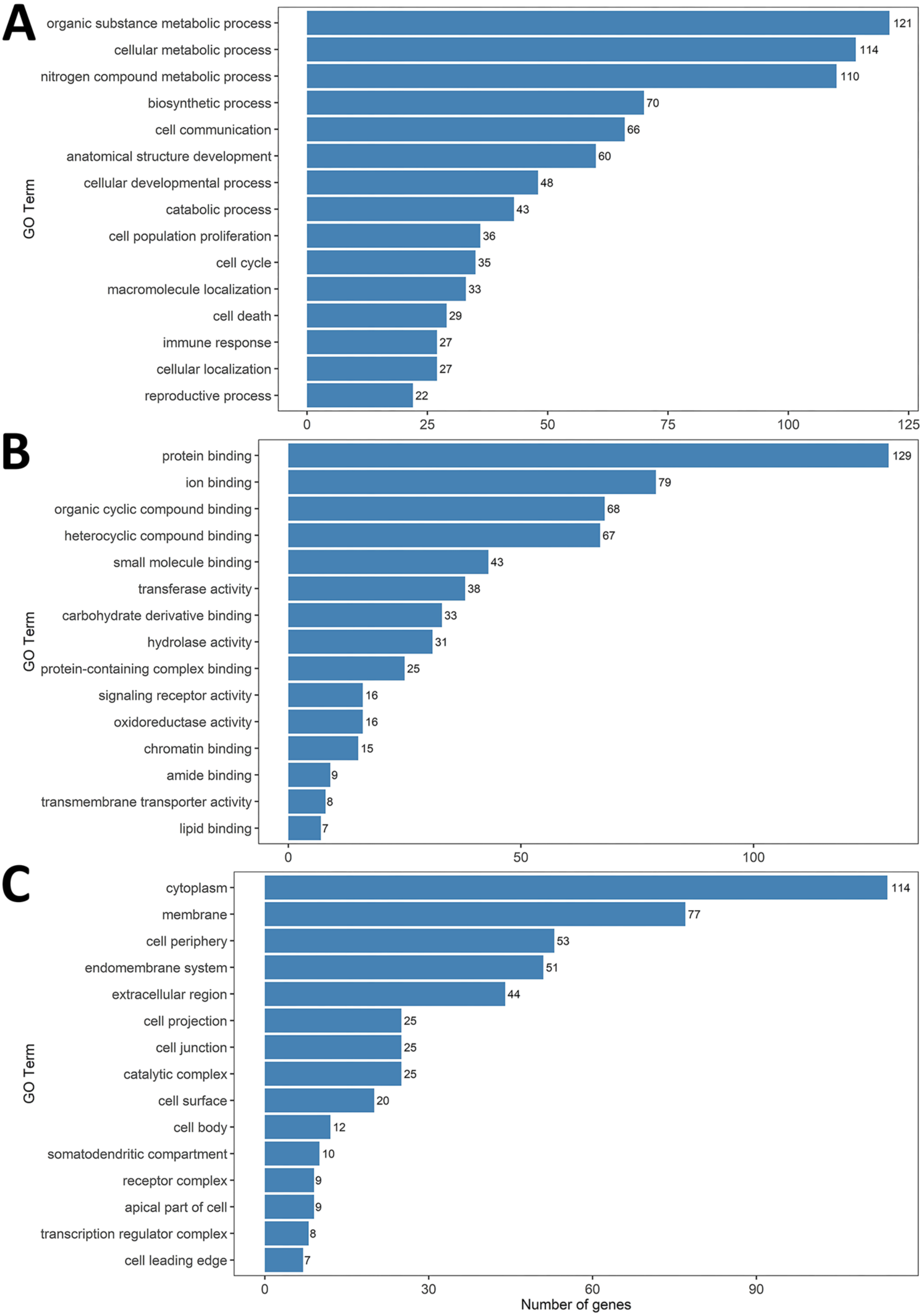


**Figure S4.** Bar plots representing the retrieved GO terms at the third level of the GO hierarchy relative to (A) Biological Processes (BP), (B) Molecular Functions (MF) and (C) Cellular Components (CC).

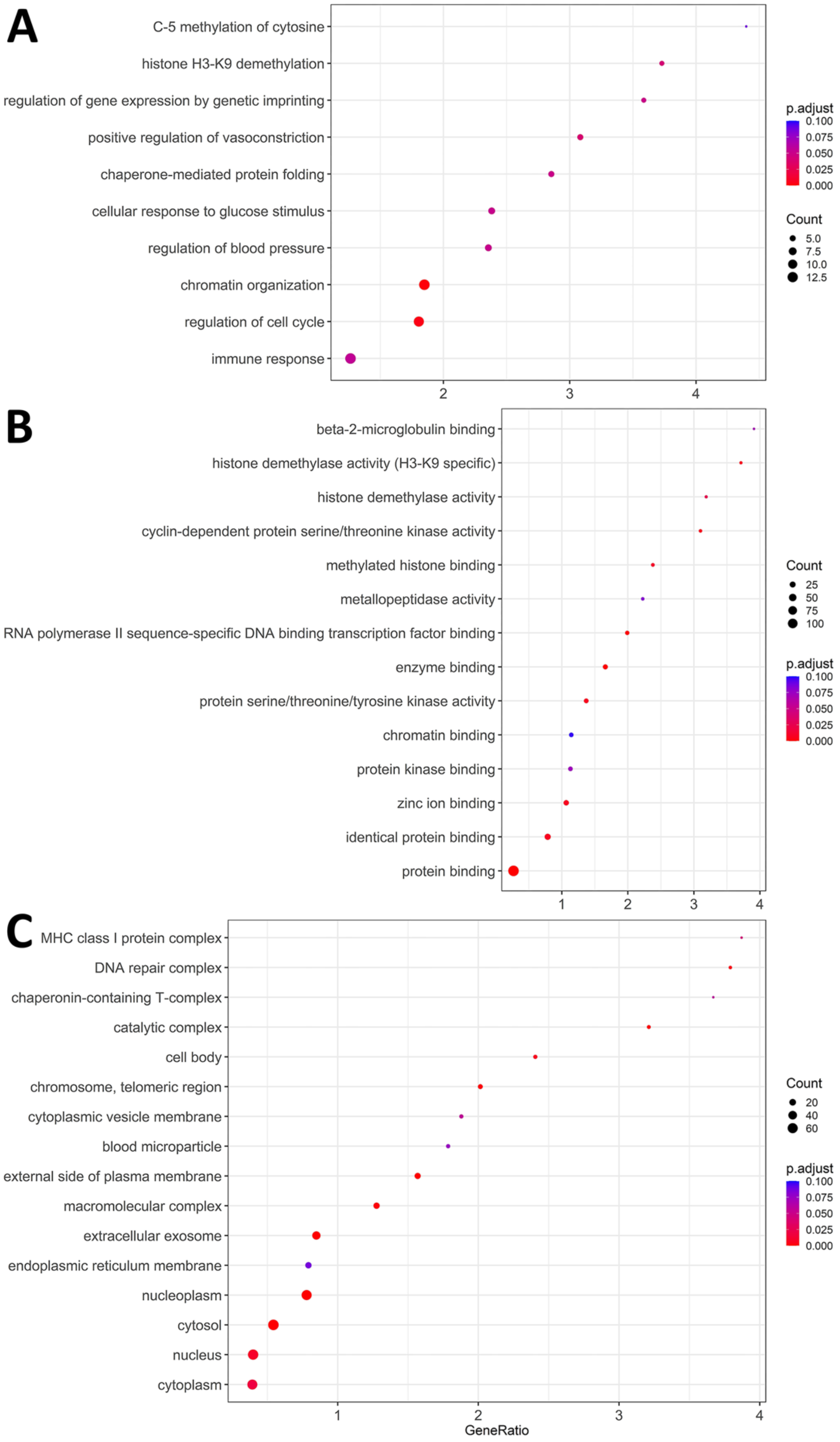


**Figure S5.** Dot plots representing the retrieved GO terms at the third level of the GO hierarchy relative to (A) Biological Processes (BP), (B) Molecular Functions (MF) and (C) Cellular Components (CC). The x-axis represents the GeneRatio, i.e. the proportion of genes in each GO term that are present in the retrieved gene list compared to the total number of genes in that GO term. The y-axis represents the statistically significant GO terms with an adjusted p-value < 0.1. The color of the dots represents the adjusted p-value (BH), red represents the smaller values, indicating higher statistical significance of the term, while blue represents larger values, indicating lower statistical significance. The size of the dots represents the number of enriched genes in the gene list associated with each GO term.

**Methods – Molecular Modelling and Dynamics**

The structure of the strychnine molecule refined using MOE at neutral pH and salt concentration of 0.15 M is reported in Figure S6.

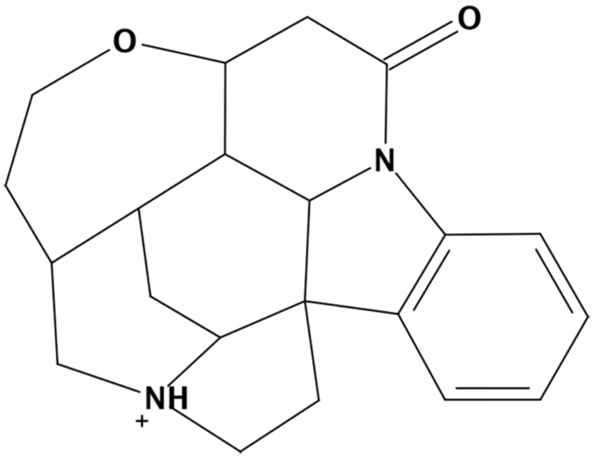


**Figure S6.** Strychnine molecule refined using MOE at neutral pH and salt concentration of 0.15 M.

**References**

1. Xu, W. *et al.* Structural basis for strychnine activation of human bitter taste receptor TAS2R46. *Science* **377**, 1298–1304 (2022).
